# Arf1-Ablation-Induced Neuronal Damage Promotes Neurodegeneration Through an NLRP3 Inflammasome–Meningeal γδ T cell–IFNγ-Reactive Astrocyte Pathway

**DOI:** 10.1101/2020.03.27.012526

**Authors:** Guohao Wang, Weiqin Yin, Hyunhee Shin, Steven X. Hou

**Author notes:** Correspondence (S.X.H.).

## Abstract

Neurodegenerative diseases are often initiated from neuronal injury or disease and propagated through neuroinflammation and immune response. However, the mechanisms by which injured neurons induce neuroinflammation and immune response that feedback to damage neurons are largely unknown. Here, we demonstrate that Arf1 ablation in adult mouse neurons resulted in activation of a reactive microglia–A1 astrocyte–C3 pathway in the hindbrain and midbrain but not in the forebrain, which caused demyelination, axon degeneration, synapse loss, and neurodegeneration. We further find that the Arf1-ablated neurons released peroxided lipids and ATP that activated an NLRP3 inflammasome in microglia to release IL-1β, which together with elevated chemokines recruited and activated γδT cells in meninges. The activated γδ T cells then secreted IFNγ that entered into parenchyma to activate the microglia–A1 astrocyte–C3 neurotoxic pathway for destroying neurons and oligodendrocytes. Finally, we show that the Arf1-reduction-induced neuroinflammation–IFNγ–gliosis pathway exists in human neurodegenerative diseases, particularly in amyotrophic lateral sclerosis and multiple sclerosis. This study illustrates perhaps the first complete mechanism of neurodegeneration in a mouse model. Our findings introduce a new paradigm in neurodegenerative research and provide new opportunities to treat neurodegenerative disorders.

## Introduction

Neurodegenerative diseases (NDs) often start with neuronal damage or disease. Alzheimer’s disease starts with extracellular aggregates formed by amyloid beta (β) peptides produced by neurons (Long and Holtzman, 2019); Parkinson’s disease starts with protein aggregates of α-synuclein from neurons; Frontotemporal dementia/amyotrophic lateral sclerosis (FTD/ALS) starts with TAR DNA-binding protein 43 (TDP-43)/Tau/Fused in sarcoma (FUS) and /or ubiquitin from neurons (Wallings et al., 2019). Therefore, studies of NDs were initially centered on neurons. In recent years, scientists began to realize the importance of immune cells in NDs. However, most studies have focused on the nonspecific, innate immune cells (mostly microglia) (Heneka et al., 2013; Hong et al., 2016; Ising et al., 2019; Ito et al., 2016; Keren-Shaul et al., 2017; Labzin et al., 2018; Singhal et al., 2014; Xu et al., 2018). More recently, a few of reports revealed T cells’ contributions to NDs (Gate et al., 2020; Jelcic et al., 2018; Lodygin et al., 2019; Shichita et al., 2009; Sulzer et al., 2017; Sun et al., 2018). Accumulated evidence suggests that the neurodegenerative process involves communication and interaction among neurons, microglia, astrocytes and immune cells (including T cells and other innate or adaptive cells). However, the mechanisms by which these cells communicate and interact to drive disease progression are largely unknown.

In this study, we investigate the mechanism by which *Arf1* knockout mediates neurodegeneration in the adult central nervous system. We find that in *Arf1*-ablated mice reactive microglia–A1 astrocyte–C3 axis was activated that promoted demyelination, axon degeneration, and synaptic loss, particularly in the hindbrain and spinal cord but not in the forebrain. We show that the neurodegenerative phenotypes of *Arf1*-ablated mice are strongly ameliorated by knockout of *IFN*γ, *Rag1*, and *NLRP3* but not of *TLR4*. The mutant phenotypes were also significantly rescued by neutralizing antibodies of VLA-4, γδ T cell receptor, and IFNγ. We further find that ablation of *Arf1* in adult neurons released peroxided lipids and ATP that activated the inflammasome in microglia to release IL-1β, which further activated γδ T cells in meninges to produce IFNγ, that entered into parenchyma to activate the microglia–A1 astrocyte–C3 neurotoxic pathway to destroy neurons and oligodendrocytes.

## Results

### Arf1 ablation induces neurodegenerative behaviors in adult mice

To determine Arf1’s function in adult mice, we generated ubiquitous deletion of Arf1 using *UBC-CreER*. After injecting *UBC-CreER/Arf1*^*f/+*^ and *UBC-CreER/Arf1*^*f/f*^ mice with tamoxifen (100 mg/kg) for five consecutive days (**Figure S1a**), we monitored their behavior. The injected *UBC-CreER/Arf1*^*f/f*^ mice (hereinafter referred to as “Arf1-ablated mice”) were normal in the first 10 days but started to show abnormal phenotypes between days 10–15. The phenotypes became more severe between weeks three and four, and the mice finally died around weeks four and five (**Figures S1b and S1c, Videos S1 and 2**). The UBC-CreER/Arf1^f/+^ control mice did not show any abnormality. The abnormal phenotypes of the Arf1-ablated mice included weight loss, reduced swimming ability, and longer time taken to walk a distance (**Figures S1b and S1c**). Later stages of the Arf1-ablated mice also displayed splaying of hindlimbs; an uncoordinated pace (**Figures S1d and S1e**); muscle atrophy (**Figure S2a**); and other abnormal phenotypes, including shivering or shaking while walking, tremors at rest, listlessness, lethargy, ataxia, and scruffy and immobility. At the last stage, they could not eat and drink and had to be euthanized. The phenotypes did not depend on normal mouse age but depended on days after Arf1 knockdown. Arf1 knockdown resulted in similar phenotypes in two-, four-, and ten-month-old mice (**Figures S1b and S1c**).

### Arf1 ablation promotes demyelination, axonal degeneration, and synapse loss

Because the abnormal phenotypes of Arf1-ablated mice are similar to those of hypomyelinating mice (Corbin et al., 1996), we examined how *Arf1* loss affects axons of the cerebellum and spinal cord of Arf1-ablated mice. From three to four weeks after tamoxifen injection, we observed increased neurological scores (**Figure S2b**), a marked reduction in the number of myelinated axons, and increased G-ratio (axon diameter/axon-plus-myelin diameter) due to demyelination (**Figures 1a–f**). Luxol fast blue staining further confirmed the demyelination phenotype (**Figures S2c and S2d**). We also examined other myelination-related proteins, including component myelin structures (Mbp, Mog, Mpz), marker of initiating myelination (CNP), and marker of mature oligodendrocytes (CC1). We found that all of these markers were significantly reduced in Arf1-ablated mice (**Figures S2e–g**). By quantifying colocalized pre- and postsynaptic puncta (synaptophysin and postsynaptic density 95 [PSD95] [**Figures 1g and 1h**]), we found a significant loss of synapses in the spinal cord and hindbrain areas (including the medulla, pons, cerebellum and midbrain) but not the forebrain areas (including the hippocampus and cerebral cortex) of Arf1-ablated mice between three and four weeks after tamoxifen injection In addition, we observed axon degeneration and denervation of neuromuscular junctions in the tibialis anterior muscle (**Figures S2h and S4a–e**) as well as neurofilament fragmentation (**Figures 1i and 1j**). Thus, Arf1 deficiency promotes demyelination, axon degeneration, synapse loss, and neurodegeneration.

**Figure 1.**
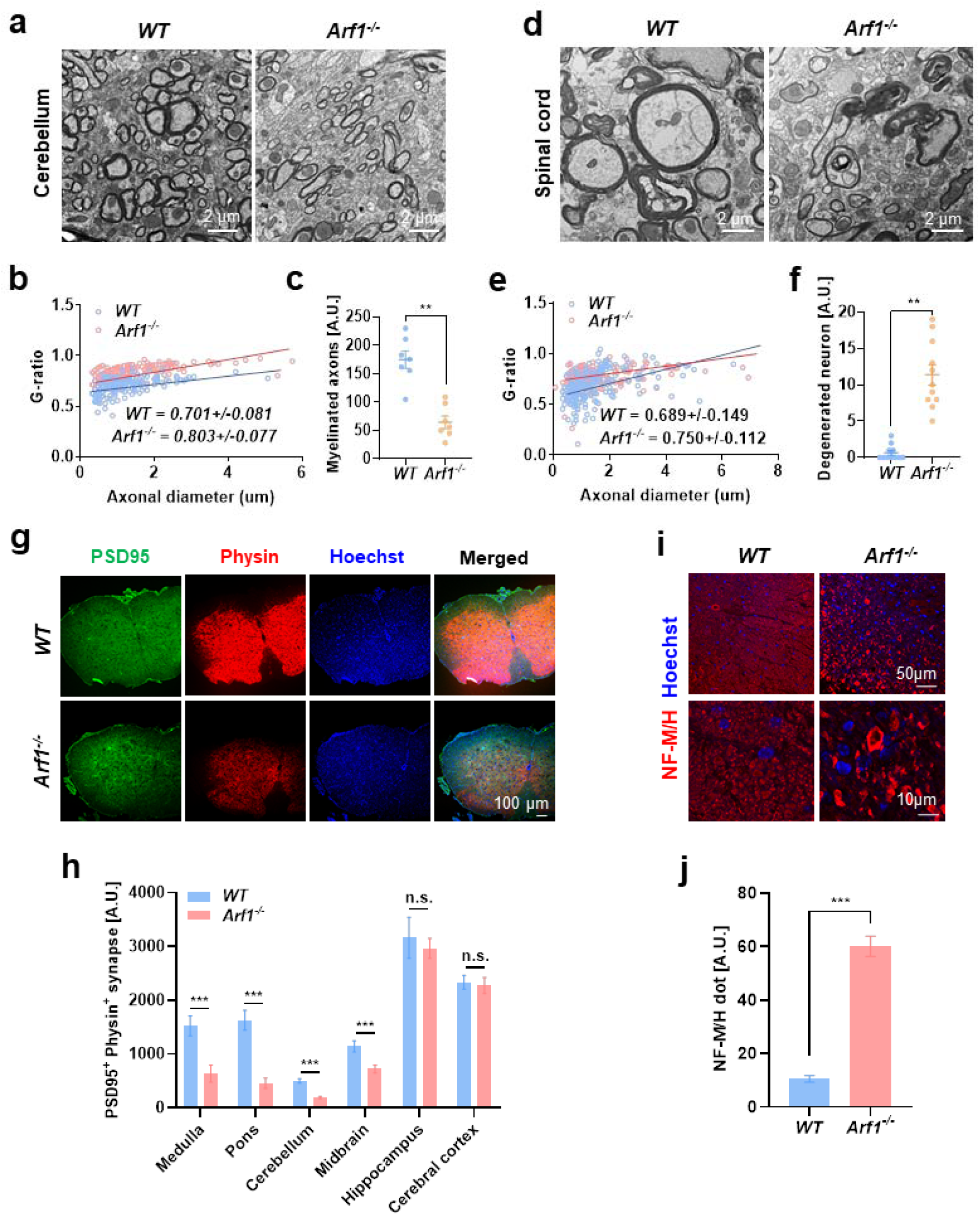
Arf1 ablation promotes demyelination, axon degeneration, synapse loss, and fragmentation of neurofilament. Control (*UBC-CreER/Arf1*^*f/+*^, WT) and Arf1-ablated (*UBC-CreER/Arf1*^*f/f*^, *Arf1*^-/-^) mice were assayed on the following indicated phenotypes. (a, b) Electron microscopy (EM) images (a) and G-ratio (b) of axons in the cerebellum of control and Arf1-ablated mice. Scale bar: 2 μm. (c) Quantification of myelinated axons in the EM sections. (d, e) EM images (d) and G-ratio (e) of axons in the spinal cords of control and Arf1-ablated mice. Scale bar: 2 μm. (f) Quantification of the numbers of degenerated neuron cells in the spinal cords of control and Arf1-ablated mice. (g) Immunofluorescence staining for PSD95 and synaptophysin in spinal cord sections from mice with the indicated genotypes. Scale bar: 100 μm. (h) Quantification of PSD95-positive and synaptophysin-positive synapses in the spinal cords of control and Arf1-ablated mice. (i) Immunofluorescence staining for NF-M/H in spinal cords from control and Arf1-ablated mice. Scale bar: 50 μm (upper layer), 10 μm (bottom layer). (j) Quantification of NF-M/H dots in panel i. n = 5 per genotype. Data are from three independent experiments and are represented as mean ± SEM. *P < 0.05, **P < 0.01, ***P < 0.001 using two-tailed t-test.

### Arf1 neuronal ablation promotes neurodegeneration

To determine the cell types involved in mediating Arf1-deficiency-induced neurodegeneration, we generated linage-specific deletion of Arf1 using *GFAP-CreER* (astrocyte; **Figure 2a**), *Pdgfra-CreER* (oligodendrocyte precursor; **Figure S3a**), *Plp1-CreER* (oligodendrocytes and Schwann cell; **Figure S3b**), *Sox10-CreER* (oligodendrocyte lineage cell; **Figure S3c**), *LysM-CreER* (myeloid cell; **Figure S3d**), *CX3CR1-CreER* (microglia; **Figure S3e**), *TMEM119-CreER* (microglia; **Figure S3f**), and *Nestin (Nes)-cre*. Among these Cre lines, only *Nes-Cre/Arf1*^*f/f*^ mice reproduced the neurodegenerative phenotypes (**Figures 2b–e, S9, and S10, Videos S3 and S4**). We further generated Arf1 deletion in neurons by using *Thy1-CreER* and reproduced most of the neurodegenerative phenotypes observed with *UBC-CreER* (**Figures 2f and S11, Videos S5 and S6**). Since *UBC-CreER* had the strongest expression and phenotypes in the brain after tamoxifen injection, we used the *UBC-CreER* line in the following experiments. These results suggest that specific ablation of *Arf1* in neurons causes the neurodegenerative phenotype.

**Figure 2.**
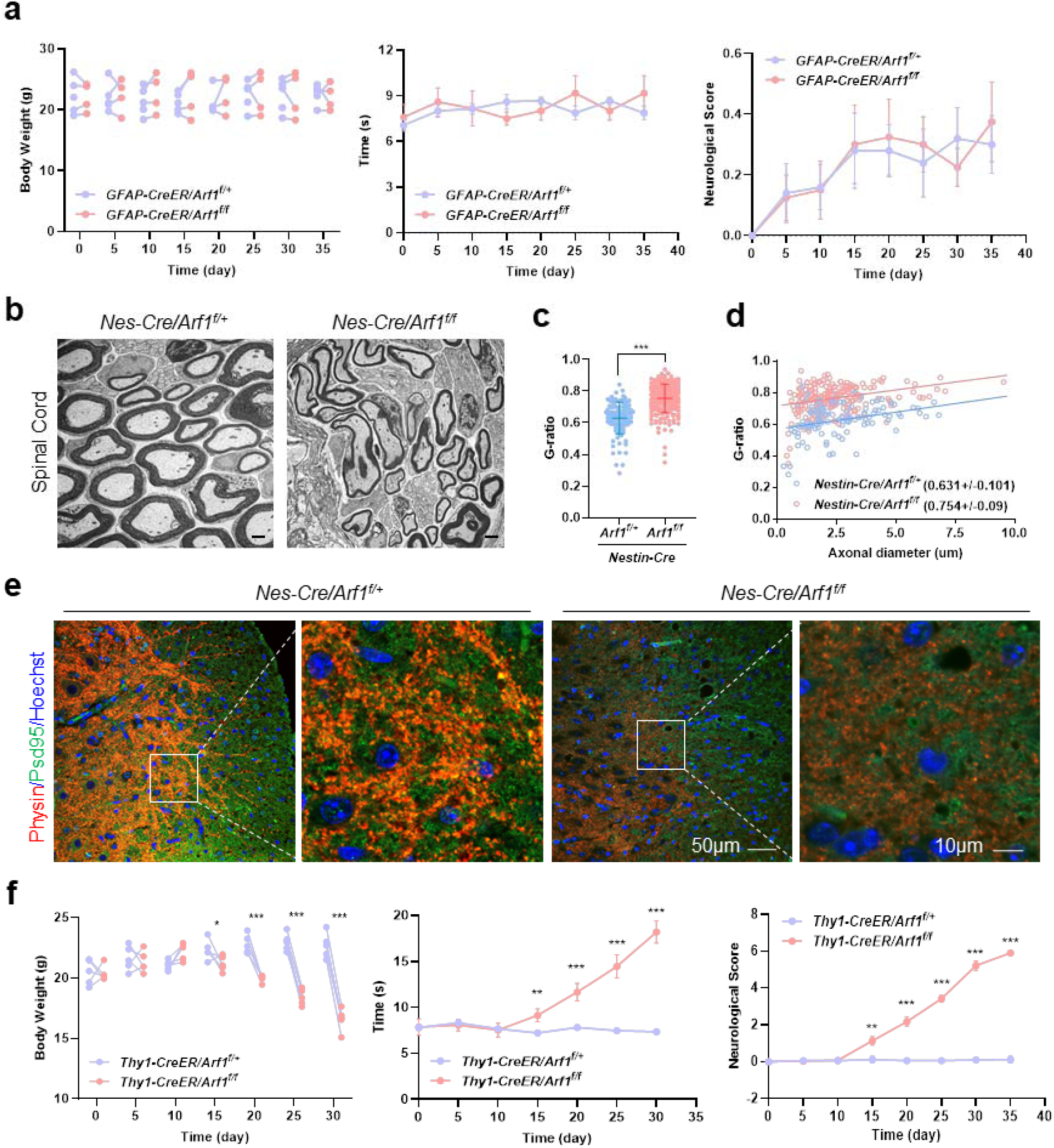
Neuronal-specific ablation of Arf1 induces the neurodegenerative phenotypes in mice. (a) Body weight, forced swimming test, and neurological score were assayed in control and astrocyte Arf1-deleted (*GFAP-CreER/Arf1*^*f/f*^) mice. (b) Electron microscopy to show axonal myelination in control (*Nes-Cre/Arf1*^*f/+*^) and neuronal Arf1-ablated (*Nestin-Cre/Arf1*^*f/f*^) mice. Scale bar: 2 μm. (c, d) Mean G-ratio and G-ratio of control (*Nes-Cre/Arf1*^*f/+*^) and neuronal Arf1-ablated (*Nes-Cre/Arf1*^*f/f*^) mice. (e) Immunofluorescence staining of PSD95-positive and synaptophysin-positive synapses in spinal cords of control (*Nes-Cre/Arf1*^*f/+*^) and neuronal Arf1-ablated (*Nes-Cre/Arf1*^*f/f*^) mice. Scale bar: 50 μm (left), 10 μm (right). (f) Body weight, forced swimming test, and neurological score were assayed in control and neuronal Arf1-ablated (*Thy1-CreER/Arf1*^*f/f*^) mice. Data are from three independent experiments and are represented as mean ± SEM. *P < 0.05, **P < 0.01, ***P < 0.001 using two-tailed t-test.

### Arf1 ablation promotes neurodegeneration by inducing IFNγ, and activating the reactive astrocyte–C3 pathway

In mouse tumors, we previously found that Arf1 ablation in cancer stem cells promoted an IFNγ-mediated antitumor immune response (Wang et al., 2020), and recent work also demonstrated that IFNγ, from meningeal T cells regulates neuronal connectivity and social behaviors (Filiano et al., 2016). To assess the potential role of IFNγ, in mediating the neurodegenerative pathway in Arf1-ablated mice, we examined the phenotypes of Arf1-ablated mice in an IFNγ-deficient background (**Figure 3**). We found that the IFNγ, protein level and activated microglia were significantly increased (**Figures 3a** and **3b**). However, the increased activated microglia were not colocalized with the lysosomal protein CD68 (**Figure S4h**), indicating that they were not activated to engulf synaptic material as previously described in Alzheimer mouse models (Hong et al., 2016). The slow traveling in the balance beam tests, poor neurological score, synapse loss, axon demyelination, and axon degeneration associated with Arf1-ablated mice were almost completely suppressed in IFNγ-deficient mice (**Figures 3c–e and S4a–g, Videos S7 and S8**). Furthermore, reactive A1 astrocytes and complement C3 were significantly increased in Arf1-ablated mice, but their increase—as well as the increase of activated microglia—were dramatically suppressed in *Arf1* and *INF*γ, double-knockout mice (**Figures 3f–i and S5**). However, the increase in activated microglia, A1 astrocytes, and C3, as well as their suppression by IFNγ, deficiency, was only restricted to the hindbrain and spinal cord, not the forebrain (**Figure S6a**). Furthermore, Stat1 phosphorylation was significantly increased in *Arf1*-deficient mice’s spinal cords, medulla, and cerebellums (**Figure S6b**). The levels of TNFα, IL-1α, and C3, but not C1q, were also significantly induced in Arf1-deficient mice but suppressed in *Arf1* and *IFN*γ, double-knockout mice (**Figures S6c–g**). However, IFNγ, deletion did not suppress increased lipid droplet (LD) accumulation and oxidized lipids in the spinal cords of Arf1-deficient mice (**Figure S7**), indicating that the lipid phenotypes are events that occur upstream of IFNγ. In addition, the slow traveling, poor neurological score, and microglial activation, but not increased oxidized lipids, were significantly suppressed by the microglial activation inhibitor minocycline (**Figures S8a–g**).

**Figure 3.**
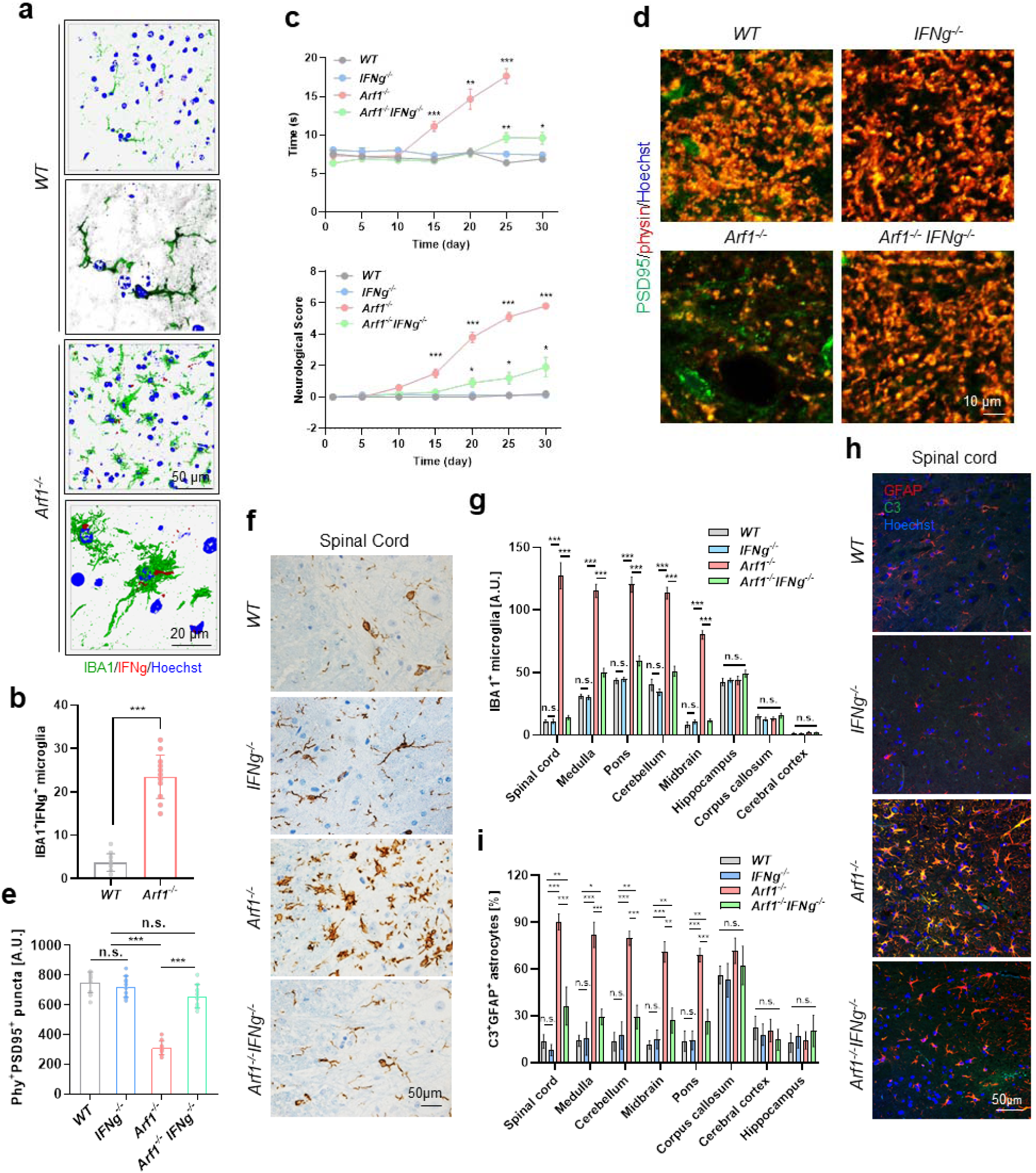
Arf1 ablation promotes neurodegeneration through induction of IFNγ, and activation of the reactive astrocyte–C3 pathway. Control (*UBC-CreER/Arf1*^*f/+*^, WT), IFNγ, deficient, (*INFg*^-/-^), Arf1-ablated (*UBC-CreER/Arf1*^*f/f*^, *Arf1*^-/-^), and Arf1-ablated in an IFNγ-deficient background *(UBC-CreER/Arf1*^*f/f*^*/INFg*^-/-^, *Arf1*^*-/-*^*INFg*^-/-^) mice were assayed on the following indicated phenotypes. (a) IFNγ, protein level and activated microglia were significantly increased in Arf1-ablated mice. Scale bar: 50 μm (upper), 10 μm (bottom). (b) Quantification of IFNγ- and IBA1-immunoreactive microglia in the spinal cords of Arf1-ablated and control mice. (c) Balance beam test and neurological score of mice with the indicated genotypes (n = 5 per group, data from three independent experiments). (d, e) Immunofluorescence staining for PSD95 and synaptophysin (d) and quantification (e) of PSD95- and synaptophysin-immunoreactive dots in spinal cord sections of mice with the indicated genotypes. Scale bar: 10 μm. (f) Immunofluorescence staining for IBA1 in spinal cords from mice with the indicated genotypes. Scale bar: 50 μm. (g) Quantification of IBA1^+^ microglia in different brain areas of mice with the indicated genotypes (n = 8 per group). (h) Immunofluorescence staining for GFAP and C3 in spinal cords from mice with the indicated genotypes. Scale bar: 50 μm. (i) Quantification of GFAP^+^ C3^+^ astrocytes in different brain areas of mice with the indicated genotypes (n = 8 per group). Note that IFNγ, deficiency suppressed increased IBA1-positive microglia and GFAP/C3-positive astrocytes of Arf1-ablated mice in areas of the hindbrain and midbrain but not the forebrain. Data are represented as mean ± SEM. *P < 0.05, **P < 0.01, ***P < 0.001 using two-tailed t-test.

We also generated *Arf1* deletion in *Tlr4* deficient mice and found that *Tlr4* deficiency did not affect the neurodegenerative phenotypes (**Figure S8h**), suggesting the neurotoxic reactive astrocytes are induced through an IFNγ–Stat1 pathway in Arf1-ablated mice, rather than by activated microglia through the TLR4–NF-κB pathway as previously reported (Liddelow et al., 2017; Yun et al., 2018).

Furthermore, the neurodegenerative phenotypes associated with Arf1 deletion by *UBC-CreER* could be reproduced in *Nes-Cre/Arf1*^f/f^ and *Thy1-CreER*/*Arf1*^f/f^ mice (**Figures 2b–f, S9, S10, and S11**), including axon demyelination, axon degeneration, synapse loss, activation of the microglia–A1 astrocyte–C3 pathway, and increased Stat1 phosphorylation. These data again suggest that the neuronal Arf1 deficiency promotes neurodegeneration.

These data collectively suggest that IFNγ, is a major downstream component in the Arf1-ablation-induced neurodegenerative pathway. IFNγ, possibly first activates microglia through the IFNγ, receptor–Stat1 signal transduction pathway (Tsuda et al., 2009) to induce expression of TNFα and IL-1α. These factors further activate A1 astrocytes to produce C3 to damage neurons and oligodendrocytes (Liddelow et al., 2017; Yun et al., 2018).

### IFNγ, comes from meningeal γδ T cells

T cells are the major source of IFNγ. We examined immune cells isolated from brain parenchyma by fluorescence-activated cell sorting analysis (**Figures S12a and S12b**). We found that T and B cells did not change, and only macrophages (CD11b^hi^F4/80^hi^) and monocytes (CD11b^hi^Ly6G/C^hi^) were significantly increased in *Arf1*-ablated mice. CD11b and F4/80 are also microglial markers, and Ly6G/C also labels microglial precursors (Getts et al., 2008). These changes might actually reflect the increased microglia as described in the earlier section. We further stained brain sections (**Figures S12c–f**) with anti-CD3 (T-cell marker), anti-Calr, anti-HMGB1, and anti-CD8α (CD8 T-cell marker) and did not find a significant change in T cells in Arf1-ablated mice compared to control mice. We also stained the brain sections with Calr and HMGB1 antibodies and found that these proteins did not significantly change between control and Arf1-ablated mice, suggesting that they are not involved in Arf1’s function in the brain.

We next used fluorescence-activated cell sorting analysis to examine immune cells isolated from brain meninges (**Figures 4a and S13a**) and found that only γδ T cells were significantly increased. To further confirm that the γδ T cells are the source of IFNγ, we generated Arf1-ablated mice in a Rag1-knockout background (*Arf1*^*-/-*^*Rag1*^*-/-*^) (**Figures 4b–g**). We found that Rag1 knockout significantly suppressed the phenotypes of Arf1-ablated mice (**Figures 4b–d and S13b–h, Video S9**). Furthermore, Rag1 knockout blocked induction of IFNγ, and C3, but not IL-1β, in Arf1-ablated mice (**Figures 4e–g**). In addition, Rag1 knockdown did not suppress the increased lipid peroxidation associated with Arf1-ablated mice (**Figures S14a and S14b**), suggesting that IL-1β production and increased peroxided lipids occur upstream of T-cell activation in Arf1-ablated mice.

**Figure 4.**
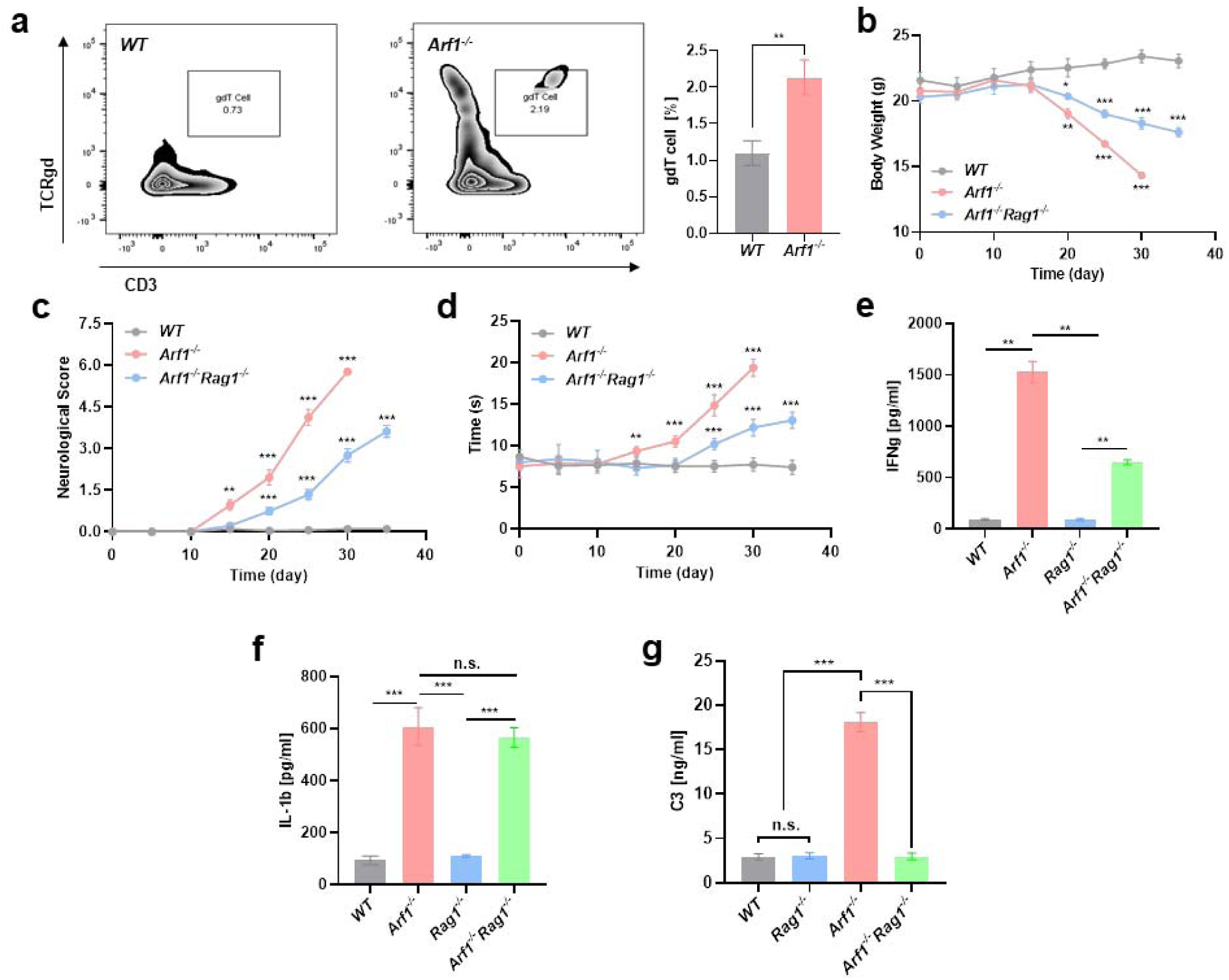
IFNγ, comes from meningeal γδ T cells. Control (*UBC-CreER/Arf1*^*f/+*^, WT), *Rag1*-deficient, (*Rag1*^-/-^), *Arf1*-ablated (*UBC-CreER/Arf1*^*f/f*^, *Arf1*^-/-^), and *Arf1*-ablated in a *Rag1*-deficient background *(UBC-CreER/Arf1*^*f/f*^*/Rag1*^-/-^, *Arf1*^*-/-*^*Rag1*^-/-^) mice were assayed on the following indicated phenotypes. (a) Meninges were dissected, and single-cell suspensions were immunostained. T cells were gated on live, single, CD3^+^, TCRγδ^+^ events and counted by flow cytometry (n = 5 mice per group). (b–d) Body weight (b), neurological score (c), and balance beam test (d) of mice with indicated genotypes (n = 5 mice per group). (e–g) Quantification of IFNγ, IL-1β, and C3 in the spinal cord lysates of mice with the indicated genotypes (n = 5 mice per group). Data are represented as mean ± SEM. **P < 0.01, ***P < 0.001 using two-tailed t-test.

We injected antibodies of IFNγ, VLA-4 (an integrin required for T-cell central nervous system homing), and γδ T-cell receptor into Arf1-ablated mice (**Figures S14c–e**). Partial elimination of IFNγ, and γδ T cells also significantly suppressed the phenotypes of Arf1-ablated mice. These data collectively suggest that the meningeal γδ T cells are the main resource of IFNγ, driving neurodegeneration in Arf1-ablated mice. Consistent with our finding, a recent publication found that γδ T cells in cerebrospinal fluid provide the early source of IFNγ, that aggravates lesions in spinal cord injury (Sun et al., 2018).

### Arf1-ablated neurons release peroxided lipids, chemokines, and ATP

We recently found that Arf1 ablation in stem cells and cancer stem cells resulted in LD accumulation, mitochondrial defects, reactive oxygen species (ROS) production, endoplasmic reticulum stress, and release of damage-associated molecule patterns (Calr, HMGB1, ATP) that activate an immune reaction to destroy tumors (Singh et al., 2016; Wang et al., 2020). Mitochondrial defects and ROS may further promote LD accumulation, lipid peroxidation, and neurodegeneration (Liu et al., 2015). We therefore examined LDs in brains of wild-type and *Arf1*-ablated mice using Oil Red O staining (**Figures S15a and S15b**) and found that, in comparison with those in brains of control mice, LDs were dramatically increased in brains of Arf1-ablated mice, particularly in the hindbrain areas and spinal cord but not in the forebrain areas.

We further assessed lipid peroxidation using the BODIPY 581/591 C11 lipid peroxidation sensor (BD-C11). Compared to control mouse brains, Arf1-ablated mice exhibited a dramatic increase in peroxided lipids in the hindbrain, midbrain, and spinal cord but not in the forebrain (**Figures 5a, 5b, S15c, and S15d**). Furthermore, ATP level was also significantly increased in the brains of Arf1-ablated mice compared to the brains of control mice (**Figure 5c**). Enzymatic activity for aconitase is a sensitive biochemical marker of ROS (Tretter and Ambrus, 2014). There was a gradual decrease of aconitase enzymatic activity in Arf1-ablated mice but not in control mice (**Figure S15e**), indicating an increased ROS in Arf1-ablated mice. In addition, we assessed expression of chemokine genes and found that several chemokines were significantly increased in Arf1-ablated mice, including CCL2, CCL4, CCL5, CCL20, CCL22, and CXCL10 (**Figure 5d**). Furthermore, our cell culture experiments suggest that CCL2, CCL5, and CCL22 were from neuronal cells, CCL4 and CXCL10 were from microglia, and CXCL10 and CCL20 were from astrocytes (**Figures S17e–g**). CXCL10 and CCL5 chemokines stimulated CD4^+^ and CD8^+^ lymphocyte migration in the intratumor and stromal compartment (Parkes et al., 2017), and CCL20 drove brain infiltration of Treg cells (Ito et al., 2019). They may have a similar role in driving γδ T cells into the brains of Arf1-ablated mice.

**Figure 5.**
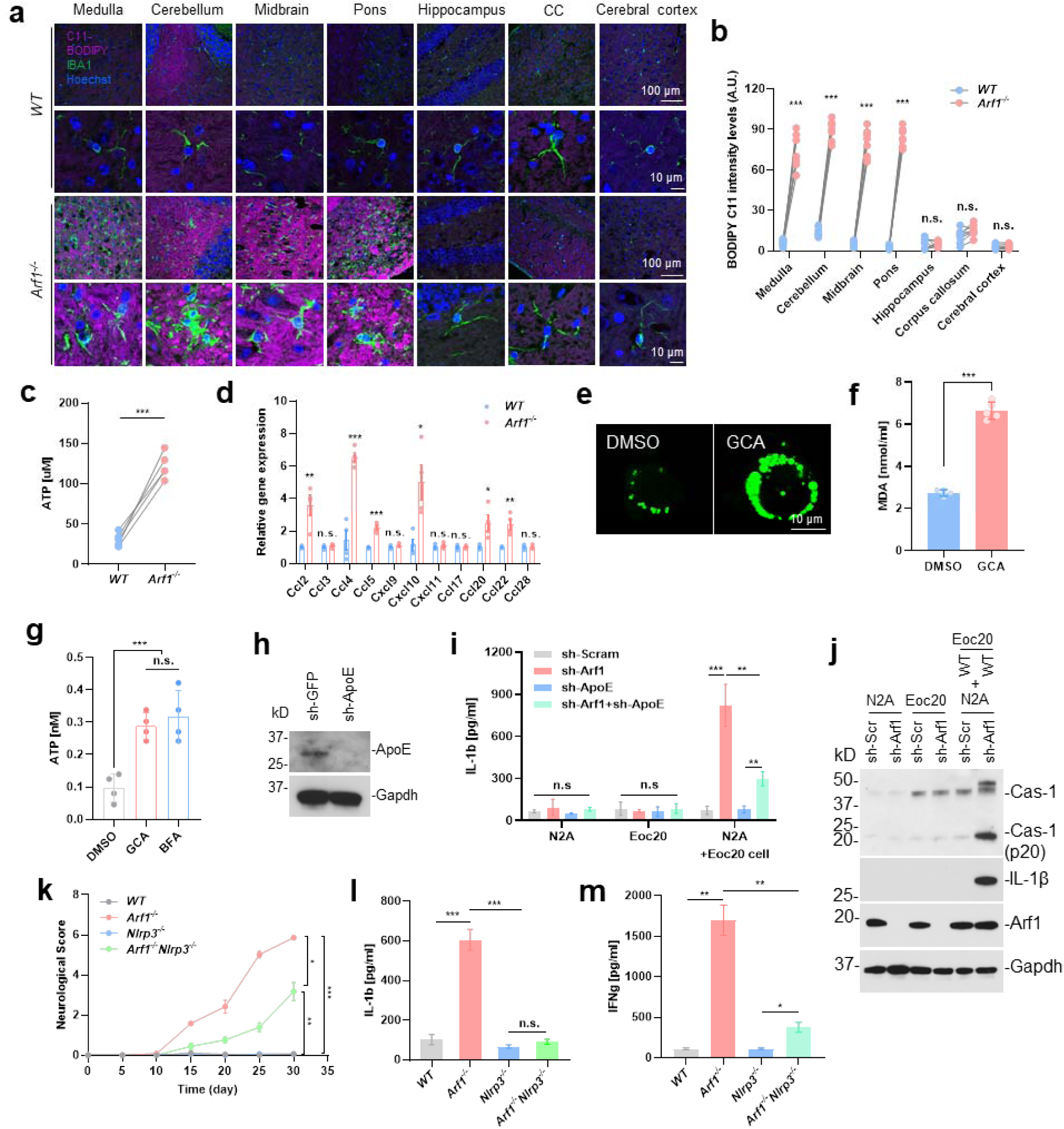
Arf1-ablated neurons release peroxided lipids, ATP, and chemokines to activate the NLRP3 inflammasome for IL-1β secretion. Control (*UBC-CreER/Arf1*^*f/+*^, WT), *Nlrp3*-deficient (*Nlrp3*^-/-^), Arf1-ablated (*UBC-CreER/Arf1*^*f/f*^, *Arf1*^-/-^), and Arf1-ablated in an *Nlrp3*-deficient background *(UBC-CreER/Arf1*^*f/f*^*/Nlrp3*^-/-^, *Arf1*^*-/-*^*Nlrp3*^-/-^) mice were assayed on the following indicated phenotypes. (a) Immunofluorescence staining for BODIPY 581/591 C11 (BD-C11), IBA1, and Hoechst in the brain sections and spinal cords from mice with the indicated genotypes. Scale bar: 100 μm (upper panel), 10 μm (bottom panel). (b) Quantification of BD-C11 level in the different brain regions and spinal cords of control and Arf1-ablated mice. (c) Quantification of ATP amount in the spinal cord lysates of mice with the indicated genotypes (n = 5 mice per group). (d) Increased chemokine levels in the spinal cord lysates of mice with the indicated genotypes. (e) BODIPY 493/503 (BD493) staining of lipid droplets in N2A cells treated with DMSO or Arf1 inhibitor GCA. Scale bar: 10 μm. (f) Malondialdehyde (MDA, a lipid peroxidation marker) in N2A cells treated with DMSO or GCA (n = 4). (g) ATP levels in culture medium of N2A cells treated with DMSO, GCA, or BFA for 24 hours (n = 5). (h) Immunoblot with anti-ApoE antibody to demonstrate that *ApoE* shRNA effectively knocked down ApoE protein expression in N2A cells. (i) Neuronal N2A cells were transfected with the indicated shRNAs, cocultured with microglial EOC 20 cells for 48 hours, and assayed for IL-1β by ELISA (n = 5 per group). (j) Western blot using antibodies of caspase-1 (Cas-1), IL-1β, and Arf1 to show that Arf1 ablation in N2A neuronal cells increased IL-1β expression and Cas-1 activation (p20) in cocultured microglial EOC 20 cells. (k–m) *Nlrp3* deficiency almost completely suppressed the neurodegenerative phenotypes of Arf1-ablated mice, including neurological score (k), induction of IL-1β (l), and induction of IFNγ, (m) (n = 5 per group). Data are represented as mean ± SEM. *P < 0.05, **P < 0.01, ***P < 0.001 using two-tailed t-test.

### Peroxided lipids and ATP activate the inflammasome in microglia to release IL-1β for activating the meningeal γδ T cells

In cultured neuron N2A cells, treatment with the Arf1 inhibitors Golgicide A (GCA) and brefeldin A (BFA) as well as knockdown by Arf1 RNAi significantly increased ATP and malondialdehyde (MDA, a lipid peroxidation marker) compared to treatment with DMSO or knockdown by Scram RNAi **(Figures 5e–g, S15f, and S15g**). In N2A and microglial cell EOC 20 co-culture, treatment of N2A cells with sh-Arf1 significantly increased incorporation of peroxided lipids into microglia (**Figures S15h-j**), and it also activated caspase-1 (p20) and increased IL-1β expression in microglia (**Figures 5h-j**). However, the increased IL-1β was significantly suppressed by knocking down *ApoE* (sh-ApoE) (**Figures 5i and S15k**), indicating that the ApoE-mediated lipid transportation is important for IL-1β induction in microglia. We also proved that the peroxided lipids MDA and 4-HNE can induce IL-1β production in microglia cells *in vitro* (**Figure S15k**). These data collectively suggest that *Arf1* knockdown in neurons promotes production of ATP and peroxided lipids, which together regulate caspase-1 activation and IL-1β production in microglia.

N-acetylcysteine amide (AD4) is an antioxidant that can penetrate the blood–brain barrier and decrease oxidative stress and LD accumulation (Ioannou et al., 2019; Liu et al., 2015). Our previous work demonstrated that Arf1 ablation promotes ATP release (Wang et al., 2020). We injected AD4 and oxidized ATP (oxATP, to block ATP function) into Arf1-ablated mice. We found that AD4 and oxATP each significantly suppressed most phenotypes associated with Arf1-ablated mice and that the two also have additive effects (**Figures S16** and **S17**). However, only AD4 suppressed the increased ROS and lipid peroxidation associated with Arf1 ablation (**Figures S16b, S16c, S16e, and S16f**).

The caspase-1 activation and IL-1β production in microglia indicate the involvement of the inflammasome in this process. To assess inflammasome involvement, we first treated Arf1-ablated mice with a specific inhibitor of the NLRP3 inflammasome (MCC950; Coll et al., 2015), which strongly suppressed phenotypes associated with Arf1 knockdown (**Figures S18a–c**). We further knocked out *Arf1* in *Nlrp3*-deficient mice and found that *Nlrp3* deficiency almost completely suppressed the neurodegenerative phenotypes of *Arf1*-ablated mice (**Figures 5k–m, S18d–l**), including induction of IL-1β (**Figure 5l**).

All the above data collectively suggest that knocking down Arf1 in neurons increases ROS and promotes the release of ATP, chemokines, and peroxided lipids; ATP and peroxided lipids activate the inflammasome to produce IL-1β in microglia; the chemokines may help to recruit γδ T cells to meninges; and the IL-1β activates γδ T cells to produce IFNγ, which enters parenchyma to activate a microglia–A1 astrocyte–C3 cascade to damage neurons and oligodendrocytes for neurodegeneration.

### Identification of a unique microglia signature associated with *Arf1*-knockout mice

To investigate the underlying molecular mechanism that regulates microglial activity in Arf1-ablated mice, we isolated microglia and used scRNA-seq-analyzed transcriptomes from wild-type control (*UBC-CreER/Arf1*^*f/+*^), *Arf1*^-/-^ (*UBC-CreER/Arf1*^*f/f*^), IFNγ-knockout (*IFNg*^-/-^) and *Arf1*^*-/-*^*IFN*γ*-/-* (*UBC-CreER/Arf1*^f/f^; *IFNg*^-/-^) mice. We performed Uniform Manifold Approximation and Projection clustering and found significant differentially expressed genes in cluster 5 of *Arf1*-ablated mice (**Figure S19a-b**), and from cluster 1 to cluster 5 microglia was identified as homeostatic (cluster 1) to disease-associated microglia (cluster 5). We also compared clusters 5 with cluster 1 in *Arf1*^*-/-*^ mice and found that cluster 5 mainly relaes to leukocyte chemotaxis and migration, tumor necrosis factor production, and regulation of inflammatory response pathway (**Figure S19c**). The cluster 5 high-expression genes include *Apoe, Cd63, Cxcl2, Cd52, Ctsb, H2-D1, Fth1*, and *Spp1* (**Figure S19d**), and cluster 5 have 22 down-regulated and 46 up-regulated genes that similar to microglia associated with neurodegenerative disease (DAM) and amyotrophic lateral sclerosis (ALS) genes (**Figure S19e, f**). The total up- and downregulation genes were shown in *Arf1*^*-/-*^ microglia compared with the other three groups (**Figures S19g and S19h**). We further compared the transcriptional profiles of Arf1-ablation-associated microglia with those of microglia in aging (**Figure S19i;** Holtman et al., 2015), ALS (**Figure S19j;** Chiu et al., 2013), DAM (**Figure S19k;** Keren-Shaul et al., 2017), Multiple sclerosis (MS) (**Figure S19l**; Kraseman et al., 2017), and found significant overlap of genes in the microglia of Arf1-ablated mice and the microglia in these disease models. The overlap includes reduced expression levels of several microglia homeostatic genes, including *P2ry12, P2ry13, Tmem119, Selplg, Cx3cr1, Hexb*, and *Siglech*, as well as upregulation of six genes: *Spp1, Ctsb, Fth1, Cd63, CD52*, and *Apoe*. Among these six, *Apoe* was one of the most upregulated. *Apoe* upregulation in microglia is a major molecular signature of NDs, and the APOE pathway plays an important role in microglial changes in NDs (Krasemann et al., 2017). In addition, the genes that are either downregulated or upregulated in Arf1-ablated microglia are more similar to genes in DAM, suggesting that the microglia in Arf1-ablated mice were activated by a Trem2-independent pathway (Keren-Shaul et al., 2017).

These data collectively suggest that the microglia in Arf1-ablated mice are a new type of DAM and that Arf1-ablated mice are a new model of NDs.

### The Arf1-ablation-induced IL-1β–γδ T cell–-IFNγ, pathway exists in MS and ALS patients

We further investigated the Arf1-downregulation-induced neurodegenerative pathway in human diseases. We first performed immunochemistry with IL-1β, IFNγ, IBA1, GFAP, and C3 antibodies on postmortem tissues from normal persons and patients with ALS and MS to examine whether increases exist in the reactive astrocyte pathway. All five markers in brains from MS and ALS patients were dramatically increased compared to those from normal persons (**Figures 6a–d**). We further examined Arf1 expression by Western blot of frozen postmortem brain stem tissues from normal persons and patients with ALS and MS. The Arf1 protein level was significantly reduced in brain tissues of ALS and MS patients compared to brain tissues of normal persons (**Figures 6e, 6f, 7a, and 7b**). The synapses also lost in the brain stem of ALS and MS patients compared with normal people (**Figures 7b, c**). These findings demonstrate that the neurodegenerative pathway associated with Arf1 downregulation exists in at least two major neurodegenerative diseases, suggesting that it may help to drive neurodegeneration. The *Arf1*-knockout mice may provide a new animal model to study ALS and MS.

**Figure 6.**
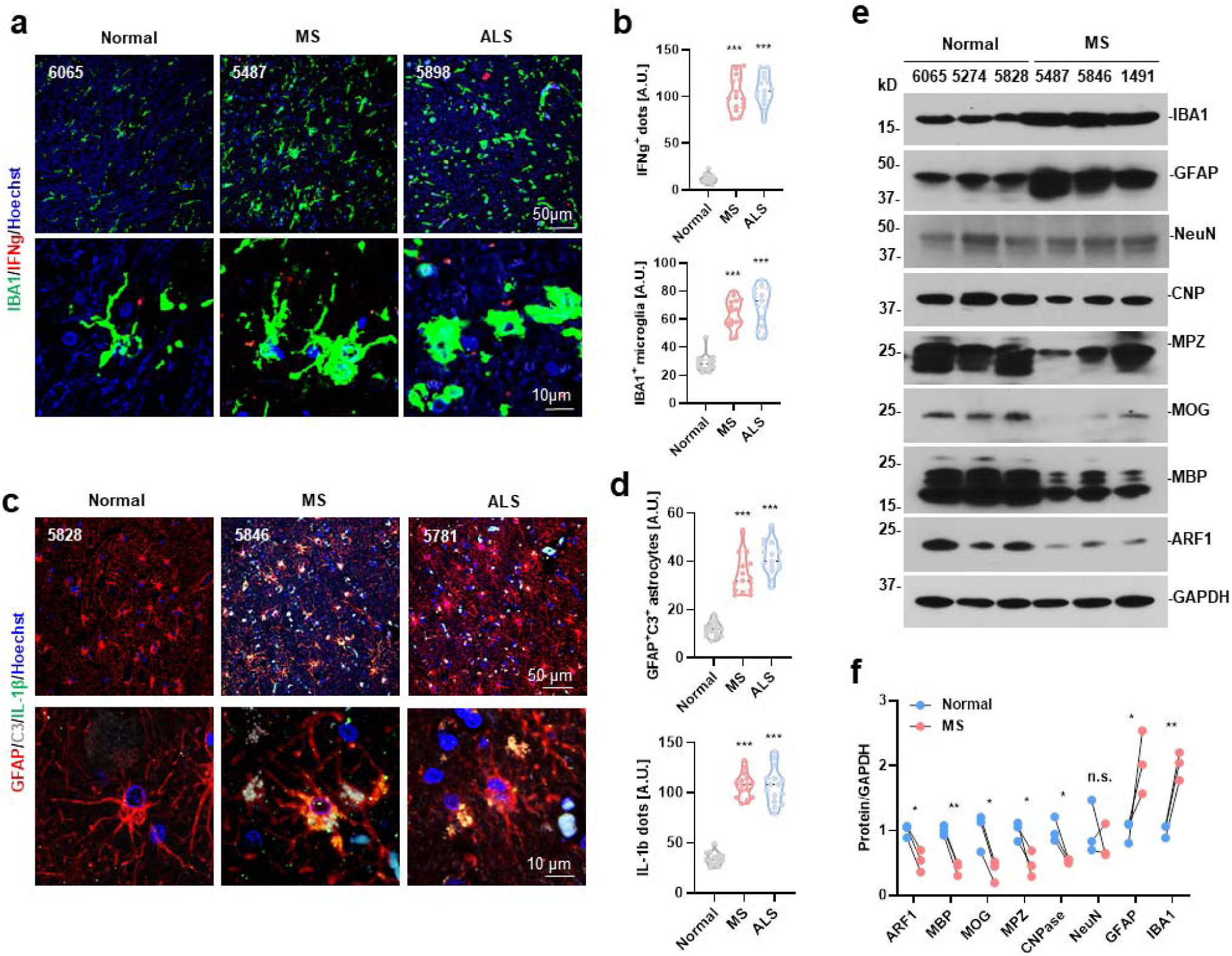
The Arf1-reduction-induced IL-1β–IFNγ–reactive astrocyte pathway exists in human diseases. (a) Immunofluorescence staining for IFNγ, IBA1, and Hoechst in pons from normal persons and MS patients. Scale bar: 50 μm (upper panel), 10 μm (lower panel). (b) Quantification of IBA1-positive microglia and IFNγ-positive dots per image field (40×, n = 12, ***P < 0.001). (c) Immunofluorescence staining for GFAP, C3, IL-1β, and Hoechst in pons from normal persons and MS patients. Scale bar: 50 μm (upper panel), 10 μm (lower panel). (d) Quantification of GFAP-positive astrocytes and IL-1β-positive dots per image field (40×, n = 12, ***P < 0.001). (e) Western blot analysis of myelin and Arf1 proteins in the spinal cords from necropsy brains of normal people and MS patients. GAPDH served as a loading control. (f) Quantitation of the ratios of proteins to GAPDH on the Western blots. Data are represented as mean ± SEM. *P < 0.05, **P < 0.01, ***P < 0.001 using two-tailed t-test.

**Figure 7.**
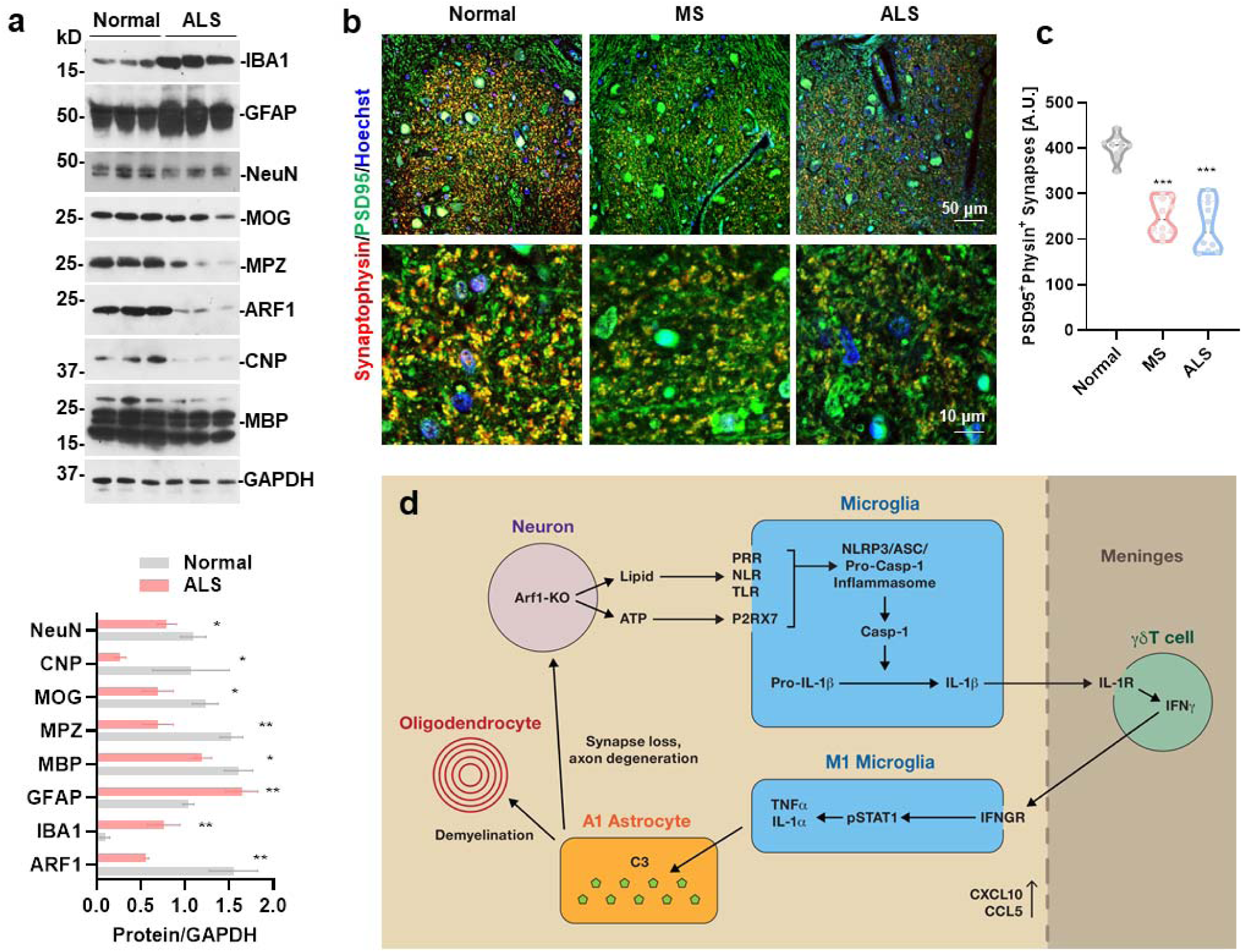
Model of the neurodegenerative pathway in Arf1-ablated mice. (a) Western blot analysis of myelin and Arf1 proteins in the medulla from necropsy brains of normal persons and ALS patients. GAPDH served as a loading control. (b) Quantitation of the ratios of proteins to GAPDH on the Western blots. (c) Immunofluorescence staining of synapse and post-synapse with anti-PSD95 and anti-synaptophysin antibodies in the medulla of control, MS, and ALS human specimens. Scale bar: 50 μm (upper panel), 10 μm (lower panel). (d) Quantification of PSD95-positive and synaptophysin-positive synapses in medulla of normal persons, MS patients, and ALS patients. (e) Proposed model of Arf1-ablation-induced neurodegeneration (see text for details). Data are represented as mean ± SEM. *P < 0.05, **P < 0.01, ***P < 0.001 using two-tailed t-test.

## Discussion

In this study, we demonstrate that Arf1 ablation in neurons results in the activation of an astrogliosis pathway (a reactive microglia–A1 astrocyte–C3 pathway) in the spinal cord and hindbrain (including spinal cord, medulla, pons, cerebellum, and midbrain) but not the forebrain (such as hippocampus, corpus callosum, and cerebral cortex). The astrogliosis pathway further causes demyelination, axon degeneration, synapse loss, and neurodegeneration. We also illustrate a comprehensive mechanism of neurodegeneration in an animal model. We find that the Arf1-ablated neurons first release peroxided lipids and ATP that activate an NLRP3 inflammasome in microglia for secreting IL-1β; the IL-1β, together with elevated chemokines, recruits and activates γδ T cells in meninges to produce IFNγ; the IFNγ, then enters into parenchyma and activates the IFNγ, receptor– Stat1 pathway in microglia to induce TNFα and IL-1α, which might further activate astrocytes to produce C3 for destruction of neurons and oligodendrocytes (**Figure 7e**).

NDs often begin with neuronal injury or disease and propagate through neuroinflammation and immune response. However, until now, the mechanisms by which injured neurons activate primary neuroinflammation and a T-cell-mediated neuropathology remained elusive. Many questions remain. First, do the neurons directly activate microglia, neuroinflammation, and T-cell immunity? In MS, one hypothesis suggests that some forms of systemic infection may initiate the whole events. The autoreactive T cells are stimulated against myelin antigens then migrate across the leaking blood–brain barrier and mediate damage against their myelin sheaths, axons, and neurons (Fletcher et al., 2010). However, there is no clear correlation between MS and infection; rather, the data indicate that, like cancer, diet and metabolism may play a more important role in MS prevalence (Yamasaki and Kira, 2019). We found that Arf1 ablation in neurons results in the release of peroxided lipids and ATP that activates the NLRP3 inflammasome in microglia to secrete IL-1β. These data suggest that neuroinflammation is initiated from inside (damaged neurons and released factors) to activate the inflammasome in microglia, rather than from outside infection.

Second, what recruits and activates T cells to the brain, and how does it do so? So far, T cells are rarely detected in brain parenchyma and mostly are restricted to the borders of the brain, particularly the choroid plexus and brain meninges and, in few cases, the cerebrospinal fluid that surrounds the brain and spinal cord due to breakdown of the blood–brain barrier (Dulken et al., 2019; Filiano et al., 2016; Gate et al., 2020; Ito et al., 2019; Keren-Shaul et al., 2017; Lodygin et al., 2019; Sun et al., 2018). It remains mostly unknown which factors trigger the recruitment of T cells to the brain. One study of Alzheimer’s disease (Gate et al., 2020) found that T cells bound two antigens produced by Epstein-Barr virus. However, no report so far has connected Epstein-Barr virus infection to neurodegeneration (Heneka, 2020). We found that several chemokines (including CCL2, CCL4, CCL5, CCL20, CCL22, and CXCL10) were significantly increased in Arf1-ablated mouse brains, which, together with peroxided lipids and secreted IL-1β from microglia, might recruit and activate γδ T cells in meninges for producing IFNγ.

Third, do the T cells kill neurons directly or indirectly? No studies have shown that T cells directly damage neurons, and only one report demonstrated that IFNγ, from γδ T cells can aggravate lesions in spinal cord injury (Sun et al., 2018). Recently, a reactive microglia–A1 astrocyte pathway has been demonstrated to mediate neuronal death during central nervous system injury and disease (Liddelow et al., 2017; Liddlow and Barres, 2017; Yun et al., 2018). Activated microglia were found to secrete a group of cytokines (IL-1α, TNF, and C1q) that induce reactive A1 astrocytes to promote the death of neurons and oligodendrocytes in various human neurodegenerative diseases, including ALS, MS, Alzheimer’s, Huntington’s, and Parkinson’s disease. A previous report found that activated astrocytes release complement C3 to damage neurons in Alzheimer’s disease (Lian et al., 2015). This information collectively suggests that a reactive microglia–A1 astrocyte–C3 pathway is possibly responsible for neuronal damage in neurodegenerative diseases. However, it is not known how this pathway is connected to T-cell-mediated neurodegeneration. We demonstrated that the reactive gliosis pathway associated with Arf1-ablated mice is strongly suppressed in IFNγ-deficient but not TLR4-deficient mice, suggesting that the pathway is activated through an IFNγ–IFNγ, receptor–Stat1 pathway in microglia (Tsuda et al., 2009), rather than by the TLR4–NF-κB pathway (Liddelow et al., 2017; Yun et al., 2018). These data demonstrate that T cells do not directly enter parenchyma through the damaged blood–brain barrier, as previously suggested; rather, they activate the reactive astrocyte pathway by releasing IFNγ. The IFNγ, connects the upstream T-cell activity to the downstream reactive gliosis pathway to damage neurons.

In addition, Arf1 is ubiquitously expressed in the brain, and its ablation in neurons only activated the reactive astrocyte pathway and caused neurodegeneration in the hindbrain and midbrain but not the forebrain. It will be interesting to find out why the forebrain is resistant to the Arf1-ablation-induced neuronal damage.

Lastly, we found that the Arf1-reduction-induced neuroinflammation–IFNγ– gliosis pathway exists in ALS and MS, suggesting that various changes described in this study may serve as biomarkers to monitor progression of neurodegeneration and that the development of new drugs that block the pathway at various points may hold great potential to treat chronic neurological diseases and acute central nervous system injuries.

## Acknowledgments

The authors thank Helen Shin for help with mouse genotyping; Kim Klarmann and Jeff Carrell at the Frederick Flow Cytometry Core for help with the flow cytometry; the Pathology/Histotechnology Laboratory at the Frederick National Laboratory for Cancer Research for help with tissue sectioning; and Kunio Nagashima at the Electron Microscopy Laboratory for help with electron microscopy experiments; Da Yin and Nathan Wong at CCR Collaborative Bioinformatics Resource, Center for Cancer Research in National Cancer Institute for help with analysis the scRNA-seq data; Dr. Samuel Lopez help for editing the manuscript. We thank the National Institutes of Health NeuroBioBank and the Rocky Mountain Multiple Sclerosis Center Tissue Bank for providing the normal, ALS, and MS human samples. This research was supported by the Intramural Research Program of the National Institutes of Health, National Cancer Institute, and Center for Cancer Research. This investigation was supported in part by a grant from the National Multiple Sclerosis Society.

## Author contributions

G.W. and S.X.H. conceived and designed the experiments. G.W. and W.Y. performed the experiments. H.S assisted with experiments. G.W. and S.X.H. analyzed the data. S.X.H. and G.W. wrote the manuscript.

## Additional information

Supplementary Information accompanies this paper.

