## supplemental information for "Arf1-Ablation-Induced Neuronal Damage Promotes Neurodegeneration Through an NLRP3 Inflammasome–Meningeal γδ T cell–IFNγ-Reactive Astrocyte Pathway"

#### Experimental Procedures

##### Antibodies, chemicals, and plasmids

Primary antibodies used in this study were purchased from the following commercial companies: PSD95 (Millipore, MAB1596), synaptophysin (Synaptic Systems, 101004), IFNγ (R&D Systems, MAB485-100), IBA1 (Thermo Fisher, PA5-27436), GFAP (Millipore, MAB360), C3 (ALZFORUM, 55444), CD3 (Cell Signaling Technology, 85061), IgG (Bio X Cell, BE0090), TCRgd (Bio X Cell, BE0070), VLA-4 (Bio X Cell, BE0071), NeuN (Millipore, M377), CNPase (Abcam, ab6319), Mpz (NOVUS Biological, NB100-1607), MAP2 (Cell Signaling Technology, 8707), Mog (Millipore, MAB5680), Mbp (Millipore, MAB386), Arf1 (Thermo Fisher, PA1-127), GAPDH (Thermo Fisher, MA5-15738), Pdgfra (BD Biosciences, 558774), CC1 (Calbiochem, OP80), Syntaxin 1 (Thermo Fisher, PA1-1042), CD68 (Bio-Rad, MCA1957GA), calreticulin (Thermo Fisher, PA3-900), HMGB1 (Thermo Fisher, PA1-16926), CD8a (Cell Signaling Technology, 85336), caspase-1 (AdipoGen, AG-20B-0042-C100), pNF-kB (Cell Signaling Technology, 3033), NF-kB (Cell Signaling Technology, 8242S), P62 (Abcam, ab109012), LC3B (Novus Biologicals, NB100-2220), caspase-3 (Cell Signaling Technology, 9662), cleaved caspase-3 (Cell Signaling Technology, 9664), Stat1 (Cell Signaling Technology, 9172S), pStat1 (Cell Signaling Technology, 8826S), PLP (Abcam, ab105784), C1q-A (Santa Cruz Biotechnology, sc-58920), anti-mouse IL-1β (R&D Systems, AF-401-NA), anti-human IL-1β (Novus Biologicals, NB600-633), anti-human IFNγ (R&D Systems, MAB285-100), anti-mouse ApoE (Novus Biologicals, NB110-60531), Olig2 (Abcam, ab109186), Neurofilament M/H (BioLegend, 801901), and LIVE/DEAD™ Fixable Aqua (Thermo Fisher, L-34965). The flow cytometry antibodies were purchased from BioLegend, as follows: anti-mouse CD16/32 (101320), CD45-APC/Cy7 (103116), CD3-Pacific Blue (100214), CD8a-AF647 (100724), CD4-PerCP/Cy5.5 (116012), TCRb-FITC (109206), TCRgd-PE (118107), NK1.1-BV650 (108735), CD25-BV785 (102051), CD127-PE/Cy7 (135013), CD11b-BV421 (101251), Ly-6G/C-FITC (108405), F4/80-BV650 (123149), CD11c-BV785 (117336), B220-PE (103208), and MHC-II-PerCP/Cy5.5 (107625). The CD1d-BV711 (740711) and Foxp3-BV421 (562996) antibodies were purchased from BD Biosciences Ltd. All secondary antibodies were purchased from Thermo Fisher Ltd.

Chemicals used in this study were purchased from the following commercial companies: BODIPY 493/503 (Thermo Fisher, D3922), BODIPY C11 (Thermo Fisher, D3861), Hoechst (Thermo Fisher, H1399), MCC950 (InvivoGen, inh-mcc), α-Bungarotoxin (Thermo Fisher, B35451), Adenosine 5'-Triphosphate (ATP) (New England Biolabs, P0756S), minocycline (Sigma-Aldrich, PHR1801), Oil Red O (Sigma-Aldrich, O0625), N-acetylcysteine amide (AD4) (R&D Systems, 5619/50), oxATP

(Sigma-Aldrich, A6779), GCA (Cayman Chemical, 18430), BFA (Thermo Fisher, B7450), MDA (MilliporeSigma, 8207560005), and 4-HNE (MilliporeSigma, 393204).

Plasmids used in this paper were purchased from the following commercial companies: sh-Scram (Addgene, 1864), sh-Arf1 (Sigma-Aldrich, TRCN0000100371, TRCN0000100373), sh-GFP (Santa Cruz Biotechnology, sc-45924-V), and sh-ApoE (Santa Cruz Biotechnology, sc-29709-V).

**Mice**

All mice were bred in the animal facility at the National Cancer Institute (NCI) at Frederick under specific pathogen-free conditions in a temperature-controlled environment with a 12-hour day/night light cycle. All breeding and maintenance was handled in accordance with the guidelines of the Animal Care and Use Committee of NCI, National Institutes of Health (NIH). The Arf1-floxed mice were generated in the animal core facility of the Mouse Cancer Genetics Program at NCI, as described previously (Wang et al., 2020). Rag1$^{-/-}$ (002216), IFNg$^{-/-}$ (002287), NLRP3$^{-/-}$ (021302), TLR4$^{-/-}$ (029015), UBC-CreER (007001), Thy1-CreER (SLICK-H) (012708), Nes-Cre (003771), Pdgfrα-CreER$^{T2}$ (032770), PLP1-CreER (005975), Sox10-iCreER$^{T2}$ (027651), GFAP-CreER (GCE) (012849), LysM-CreER$^{T2}$ (031674), TMEM119-CreER (031820), and CX3CR1-CreER (021160) mice were obtained from The Jackson Laboratory. To generate Arf1-knockout mice in different cell types, we crossed homozygous floxed Arf1 mice with different CreER transgenic mice that express tamoxifen-inducible Cre at the whole-body level or in specific cell types in the nervous system or monocytes. Double-knockout of *Arf1* with *IFNg*, *Rag1*, *NLRP3*, or *TLR4* was generated by crossing *UBC-CreER/Arf1*$^{f/f}$ mice with IFNg$^{-/-}$, Rag1$^{-/-}$, NLRP3$^{-/-}$, or TLR4$^{-/-}$ mice for three generations to get the homozygous mice. Primers were used for genotyping mice as provided by The Jackson Laboratory website. Arf1 genotyping primers were used as follows: Arf1-gP5, 5’-GGTTTTAAGAGGCCCTGTGTC-3’; Arf1-gP3’, 5’-TCGGGAGCTGGCACTAAAAA-3’; and Arf1-gR, 5’- TGCACACACCAAGTACAAGC-3’.

**Injecting Chemicals into mouse**

Tamoxifen (T5648) was dissolved in corn oil (Sigma-Aldrich, C8267) and intraperitoneally (i.p.) injected into mice at 0.1 mg/g of body weight for five consecutive days. For treatment of mice with minocycline, MCC950, AD4, and oxATP, minocycline was dissolved in saline and injected i.p. at 50 mg/kg each day for two consecutive weeks; the MCC950 was dissolved in phosphate-buffered saline (PBS), and the mice were injected i.p. with MCC950 (10 mg/kg) or PBS once every two days for two consecutive weeks; AD4 and oxATP were dissolved in PBS, and AD4 (100 mg/kg) and oxATP (15 mg/kg) were administered to each mouse by i.p. injection at 24-hour intervals for two consecutive weeks, beginning 24 hours after mice were challenged with tamoxifen five times.

**Mouse behavior test**

Mouse body weight was measured daily by using a balance after tamoxifen injection. Fine motor coordination and balance were assayed by balance beam test. Mice were trained for two days to walk the entire length of a 6-mm- or 12-mm-wide × 80-cm-long wooden beam suspended 50 cm above the floor. The time for a mouse to cross the beam was recorded. In each test, the mice were placed onto one end of the beam and ran down the entire beam into a dark box. Each genotype used five mice, and each mouse was tested three times. The results were the average of 15 trials.

For forced swimming, mice were placed separately into a 15 cm (diameter) × 20 cm (high) plastic cylinder that was filled with 12 cm of 25 °C water. Mouse immobility time was defined as absence of active behaviors, such as struggling or swimming to escape. If a mouse moved its feet and kept its head above water, it was defined to have mobility. A trained observer measured each mouse for 180 seconds.

The footprinting assay was performed as follows: black ink was applied to the hind paws and red ink was applied to the forepaws of tested mice. The mice were allowed to walk through a narrow space with white paper placed at the bottom. Stride lengths were calculated by measuring with a ruler to identify the distances between the forelegs and hind legs in each step.

Neurological score was observed daily with disease development after tamoxifen injection. Throughout disease progression, mice were scored to indicate neurological damage, as follows: 0 – no clinical signs; 1 – limp tail or hindlimb weakness; 2 – limp tail and hindlimb weakness; 3 – loss of coordinated movements; 4 – hindlimb paralysis; 5 – hindlimb and forelimb paralysis; 6 – moribund.

**Electron microscopy**

For electron microscopy, adult mice were euthanized with $CO_2$ for 30 minutes, and the brain and spinal cord were isolated using surgical scissors and tweezers. The brain and spinal cord were perfused with electron microscopy buffer containing 4% paraformaldehyde (PFA), 0.2% glutaraldehyde, and 0.1 M sodium cacodylate for 48 hours. Fixed tissues were sent to the NCI Electron Microscopy Laboratory at the Advanced Technology Research Facility in Frederick, Maryland, for sectioning and imaging.

**Western blot**

Mouse brains, spinal cords tissues, or cells were homogenized in RIPA lysis buffer (0.1% SDS, 150 mM NaCl, 0.5% sodium deoxycholate, 1 mM EDTA, 1 mM EGTA, 1.0% Triton X-100, and 50 mM Tris pH 8.0) with 1 × phosphatase inhibitor cocktail (Sigma-Aldrich) and 1 × protease inhibitor cocktail (Promega) 30 times and lysed on ice for 30 minutes, sonicated at 20 W for 30 seconds, and then centrifuged at 12,000 × *g* at 4 °C for 2 minutes to delete pellets. The saved supernatants were used for gel analysis. The protein concentration was determined by a Bio-Rad protein assay kit (Bio-Rad). A total of 300 μl of protein (2.50 μg/μl) was added with 1 × SDS loading buffer and boiled by a 100 °C heater for 15 minutes. Equal volumes of protein samples were loaded onto 4–20% Mini-PROTEAN® TGX™ precast protein gels (Bio-Rad) and run by a 120 V PowerPac™ Basic power supply (Bio-Rad) to electrophorese for 65 minutes. The gel was transferred on nitrocellulose membrane (GE Healthcare). After transfer, the membrane was blocked with 5.0% blotting-grade blocker (Bio-Rad) for 1.5 hours, and then incubated with primary antibodies at 4 °C with shaking overnight. The next day, the membrane was washed three times with 1 × PBST for 5 minutes each and incubated with the secondary HRP antibodies (1:2000) in 5% milk for 2 hours. The membrane was then developed by Pierce™ ECL Western Blotting Substrate (Thermo Fisher, 32106), applied to HyBlot CL® autoradiography film (Thomas Scientific, 3022), and developed with an S&W Imaging system (Quantum Medical Enterprises LNC.).

**RNA isolation and qRT-PCR**

Total RNA of mouse brains or spinal cords was extracted with the RNeasy Micro Kit (Qiagen, 74104), and cDNA was synthesized by the High-Capacity cDNA Reverse Transcription Kit (Applied Biosystems, 4368814). RNA concentration was determined by a NanoDrop (DS-11 spectrophotometer from DeNovix, Inc.), and an equal amount of RNA was used for cDNA synthesis. The cDNA amplification was performed with a SYBR™ Select Master Mix (Applied Biosystems, 4472903) using a CFX96 Touch™ Real-Time PCR Detection System (Bio-Rad). The *Gapdh* gene was used as an internal control. The results are shown as $2^{-\Delta CT}$. Each sample underwent three independent experiments. The pan-astrocyte, A1 astrocyte, and A2 astrocyte primer sequences were used as described previously (Yun et al., 2018). The primers used in this study are shown in Table 1.

**Cell culture**

Mice Neuro-2a (ATCC® CCL-131™) and LADMAC (ATCC® CRL-2420™) cells were purchased from American Type Culture Collection (ATCC) and cultured in Eagle's minimum essential medium supplemented with 10% fetal bovine serum (FBS), 100 U/mL penicillin, and 100 µg/mL streptomycin. HEK 293 cells were cultured in Dulbecco's modified Eagle's medium (DMEM) supplemented with 100 µg/mL streptomycin, 100 U/mL penicillin, and 10% FBS. EOC 20 (ATCC® CRL-2469™) mouse microglia cells were purchased from ATCC and cultured in DMEM containing 4 mM L-glutamine adjusted to contain 1.5 g/L sodium bicarbonate and 4.5 g/L glucose, with 10% FBS and 20% LADMAC conditioned media (produced from the LADMAC cell line [CRL-2420]).

**Immunohistochemistry and Immunofluorescence**

Mice were anesthetized by Isothesia isoflurane (Henry Schein, NDC11695-6776-2) perfused with 20 mL saline and further perfused with 4% PFA in 0.01 M PB. Brains and spinal cords were isolated and fixed by 4% PFA overnight and transferred into 30% sucrose at 4 °C to let the tissues completely sink to the bottom of the tube. The tissues were sectioned with paraffin embedding at 10 µm or frozen sectioning at 20 µm. The sections were deparaffinized with xylene and alcohol, and antigen retrieval was performed by citrate buffer (Abcam, ab93678) or Tris-EDTA buffer (Abcam, ab93684) in a 100 °C cooker for 30 minutes. Mouse brain and spinal cord slides or cultured cells were fixed with 4% PFA at room temperature for 15 minutes, washed three times with 1 × PBST, blocked with 10% goat serum for 1 hour, and incubated overnight with primary antibodies in 3% bovine serum albumin at 4 °C. The next day, the slices were washed three times with 1 × PBS and incubated with Hoechst and secondary antibodies (AF-488, 594, or 647; Thermo Fisher) for 2 hours at room temperature. The cells or sections were examined by using a Zeiss (LSM780) confocal microscope system and analyzed with Zeiss black software.

**Oil Red O, Luxol Fast Blue & Masson's Trichrome Staining**

Kits for Oil Red O, Luxol fast blue, and Masson's trichrome staining were purchased from Abcam Ltd. (Oil Red O Stain Kit [Lipid Stain, ab150678], Luxol fast blue staining kit [ab150675], and Trichrome Stain Kit [Connective Tissue Stain, ab150686]), and staining was carried out according to the instructions..

**Isolation of microglia cells from adult mice**

Mice were euthanized by $CO_2$ for 20 minutes, after which the brain and spinal cord were quickly dissected. The brain and spinal cord were transferred to 2 mL Hanks' balanced salt solution (HBSS) plus 1 × penicillin/streptomycin, after which they were minced as finely as possible using a scalpel blade. The tissue was transferred to a 15-mL tube containing 3 mL of dissociation medium (1 mg/mL papain, 1.2 U/mL dispase II, and 20 U/mL DNase I dissolved in DMEM/F12 medium). The tissue suspension was gently rocked in a tissue culture incubator for 20 minutes, then 5 mL neutralization medium was added to neutralize the enzymes. The mixture was centrifuged for 5 minutes at 250 × *g* at 25° C, the medium was aspirated slowly, and the pellets were washed twice with 5 mL of serum-free medium (4.9 mL DMEM/F12 plus 0.05 mL of penicillin/streptomycin [100×] and 0.05 mL of 0.45 g/mL glucose). They were next centrifuged for 5 minutes at 350 × *g*. The pellets were kept, 3 mL DMEM/F12 was added, and a polished Pasteur pipette with a large hole was moved up and down against the bottom of tube to break up the large clumps of tissue. The mixture was allowed to sit for 1 minute until the large clumps settled. Next, the supernatant was transferred to a 15-mL tube, another 3 mL of DMEM/F12 was added to the pellets, and a pipette was moved up and down again to transfer the supernatant to 15-mL tube. The cells were filtered with a 40-μm cell strainer, spin at 350 × *g* for 4 minutes, washed with 5 mL DMEM/F12, and then centrifuged at 350 × *g* for 4 minutes to get the pellet. The cell pellet was resuspended with 4 mL of 37% Percoll. The 15-mL tube was prepared, 4 mL of 70% Percoll was added at the bottom, 4 mL of the cell pellet in 37% Percoll was added next, another 4 mL of 30% Percoll was added on top of that, and 2 mL of HBSS was added on the top. The cells were centrifuged in the tube for 40 minutes at 300 × *g* (18 °C) with no brake. After centrifugation, the layer of debris was gently removed and 2.0–2.5 mL of the 37–70% interphase was collected into a clean 15-mL corning tube. The cells were washed twice with 6 mL HBSS and centrifuged for 7 minutes at 500 × *g* at 4 °C. The cells were sorted by CD11b beads at first, and then sorted by CSF1R beads. The sorted cells were mainly microglia. The cells were kept in ice and sent to the Advanced Technology Research Facility in Frederick, Maryland, for single-cell sequencing.

**Single-cell sequencing**

Microglia cells were sequenced by four 10x Genomics Chromium Single Cell platform. The barcoded libraries were generated following the manufacturer's specifications, and 3' mRNA libraries were made and sequenced on one NextSeq run and one NovaSeq SP run. All samples have sequencing yields of more than 315 million reads per sample. The sequencing was set up as a 28 cycles + 55 cycles non-symmetric run on the NextSeq and as a 28 cycles + 75 cycles non-symmetric run on the NovaSeq. Demultiplexing was done, allowing one mismatch in the barcodes. Over 97.2% of bases in the barcode regions have Q30 or above, at least 95.0% of bases in the RNA read have Q30 or above, and 95.6% or more of bases in the sample index have Q30 or above. More than 97.2% of bases in the unique molecular identifier (UMI) have Q30 or above.

**Analysis of single-cell RNA-seq**

The analysis was performed by the Center for Cancer Research Collaborative Bioinformatics Resource at NCI using the default parameters and aligned to the GRCm38 (mm10) mouse reference genome. The number of captured cells ranges from 27,930–30,710, and mean reads per cell ranged from 10,611–13,205. Cells with an extremely low number of UMI counts were filtered out. Median genes found per cell ranged from 1,111–1,187, and the total number of genes detected ranged from 23,003–23,633. Morethan 50% cells were recognized as microglia. Uniform Manifold Approximation and Projection (UMAP) were generated to create a map of merged cells from four samples of mice brain. Tabula Muris was used to annotate cell identities. Microglia were extracted and retransformed for all subsequent analysis. Microglias were separated into 12 clusters. Model-based analysis of single cell transcriptomics (MAST) was used to compute differentially expressed genes for each cluster compared to the rest of the clusters. Genes with an adjusted p value of 0.05 or below and an absolute log2 fold change of 0.5 or above is annotated as significant. Chi-squared test was used test for any disproportional distribution of cell numbers in each cluster. Signed value is calculated for each gene as Signed.value = sign (logFC) * log10 (P-value) * abs (log FC). Genes that fall on the top right quadrant indicates concordant upregulated with previously identified disease state genes. Genes that fall on the bottom left quadrant indicates concordant downregulated with previously identified disease state genes. DEGs from both Arf1-WT contrast and Cluster5-ClusterAll contrast showed a high number of genes that showed similar signed value signatures as those disease state genes. We computed Cluster5-cluster1 differentially expressed genes and performed the over-representation analysis using the Gene Ontology – Biological Process gene set.

**Flow cytometry analysis**

Mice meninges and brain were isolated as described previously (Manglani et al., 2018). Cell surface markers were stained with relative antibodies (purchased from BioLegend and BD Biosciences Ltd.) for 30 minutes at 4 °C in the dark. (The details of the methods performed are described on the BioLegend website.) For staining of the intranuclear transcription factor Foxp3, cell were stained with surface marker and fixed with 0.5 mL/tube Fixation Buffer (BioLegend, 422101) in the dark for 20 minutes at room temperature according to the manufacturer's instructions. Then, cells were washed twice with 1 mL of Intracellular Staining Permeabilization Wash Buffer and centrifuged at 350 x *g* for 5 minutes. The supernatant was discarded, and the cells were stained with intercellular antibody. Stained cells were analyzed with a BD LSRFortessa cell analyzer, and the data were analyzed with FlowJo 10 software.

**Enzyme-linked immunosorbent assays (ELISA)**

The mouse extracellular ATP, IL-1α, IL-1β, IFNγ, TNFα, C1Q, C3, aconitase, and MDA analyses were performed with ELISA kits purchased from commercial vendors, including the ENLITEN® ATP Assay System (Promega, FF2000), ELISA MAX™ Deluxe Set mouse IL-1α (BioLegend, 433404), Mouse IL-1β ELISA Kit (BioLegend, 432601), Mouse IFNγ ELISA Kit (BioLegend, 430801), TNF alpha Mouse ELISA Kit (Invitrogen, BMS607-3), Mouse Complement 1q (C1q) ELISA Kit (MyBioSource, MBS2702391), Mouse Complement C3 ELISA Kit (MyBioSource, MBS763294), Aconitase Assay Kit (Abcam, ab83459), and Lipid Peroxidation (MDA) Assay Kit (Colorimetric/Fluorometric) (Abcam, ab118970). Detailed methods for performing the analysis are described in the manufacturers' instructions.

**Treatment Mice with Neutralizing Antibody**

Rat IgG1 isotype control (BP0088) and neutralizing antibodies of IFNγ (BP0055), VLA-4 (BE0071), TCRg/d (BE0070), and Armenian hamster IgG (BE0091) were purchased from Bio X Cell Ltd. Each mouse was i.p. injected with neutralizing antibody of IFNγ (250 μg/mouse), TCRg/d (400 μg/mouse), and VLA-4 (300 μg/mouse) or relative amounts of control antibodies every two days for three weeks. The mice injected with control antibodies and PBS had similar phenotypes.

**Human samples**

Human postmortem brain and spinal cord tissue samples were obtained from the NIH NeuroBiobank, the University of Maryland Brain and Tissue Bank, and the Rocky Mountain Multiple Sclerosis Center Tissue Bank. All experimental procedures followed the NIH NeuroBiobank's guidelines and restrictions. Age-matched control human tissues were used for comparison with MS and ALS patients' tissues. Each group used three independent human tissues. Human sample information is described in Supplemental Table 2.

**Statistical analysis**

Each group in the mouse behavior study used at least five mice. The number of mice used is shown in the figure legend; the cell experiments were performed with triplicate samples and in two independent experiments. Data are shown as mean ± SEM or mean ± SD. Comparisons of two groups were done by paired two-tailed Student's t-test. Analysis of more than two groups was performed by one-way ANOVA (with Tukey's multiple-comparison post-tests). Statistical analysis was done with GraphPad Prism 8. A P value $< 0.05$ was considered significant.

### Supplemental Figure

**Figure S1. Arf1-deficient mice have movement defects.**

(a) Schematic diagram of the experimental strategy used to generate Arf1-knockout (ablated) adult mice. (b, c) Survival curve, body weight, forced swimming test, and balance beam test of two-month-old (b) and ten-month-old (c) adult mice. (d) Gait of control (*UBC-CreER/Arf1*$^{f/+}$, WT) and Arf1-ablated (*UBC-CreER/Arf1*$^{f/f}$, *Arf1*$^{-/-}$) mice was examined by footprinting assay at two months, four months, and ten months of age. (e) Quantification of stride lengths of front and hind footprints of mice in the indicated ages and genotypes. Data are from three independent experiments and are represented as means ± SEM. *P < 0.05, **P < 0.01, ***P < 0.001 using two-tailed t-test.

**Figure S2. Muscle atrophy, demyelination and neuromuscular denervation of Arf1-ablated mice.**

(a) Masson's trichrome stained sections of triangularis sterni (TS) muscles from 2-month-old control and Arf1-ablated mice. Scale bar: 100 µm. (b) Neurological scores of Arf1-ablated mice in comparison with controls. (n=5 mice per group). (c, d) Luxol Fast Blue Staining of cerebellum (b) and spinal cord (c) of control and Arf1-ablated mice. Scale bar as indicated. (e) Immunofluorescence staining for MBP, CD3, CNP and Hoechst of spinal cord from wild-type and Arf1-ablated mice. Scale bar: 10 µm. (f) Immunofluorescence staining for Pdgfra, CC1 and Hoechst of cerebellum from wild-type and Arf1-ablated mice. Scale bar: 50 µm (left), 10 µm (right). (g) Immunoblot analysis of Arf1, myelin proteins and Neuron N (NeuN) in spinal cords of 2-, 4- and 10-month-old control and Arf1-ablated mice. (h) Syntaxin and α-bungarotoxin immunostaining of TS muscles from 2-month-old control and Arf1-ablated mice. Scale bar: 50 µm. Data are represented as mean ± SD. **P < 0.01, ***P < 0.001 using two-tailed t-test.

**Figure S3. Ablation of Arf1 in oligodendrocytes, myeloid cells and microglia did not show neurodegenerative phenotypes.**

(a-f) Body weight, balance beam test and neurological score were assayed in control and cell-type specifically Arf1-deleted mice. a-Pdgfra-CreER (oligodendrocyte precursor); b-PLP1-CreER (oligodendrocytes and schwann cell); c-Sox10-CreER (oligodendrocyte lineage cell); d-LysM-CreER (myeloid cells); e-CX3CR1-CreER (microglia); f-TMEM119-CreER (microglia). n=5 per group, Data are represented as mean ± SEM.

**Figure S4. Arf1 ablation promotes neurodegeneration through INFγ.**

(a-e) Axon degeneration phenotypes associated with Arf1-ablated mice were strongly suppressed in INFγ deficient background. Mean axonal numbers (a), mean axonal diameters (b), individual G-ratio distribution (c), mean G-ratios (d), and distributions of axonal diameters (e) in the ventrolateral lumbar spinal cord white matters of WT, $Arf1^{-/-}$, $IFNg^{-/-}$, and $Arf1^{-/-}IFNg^{-/-}$ mice as indicated. Data are shown as Mean ± SEM. $^{**}P < 0.01$, $^{***}P < 0.001$ using two-tailed t-test. (f) Immunofluorescence staining for Mbp, Mog and Hoechst revealed that IFNγ deficiency suppressed reduction of myelin proteins in the spinal cord white matters of Arf1-ablated mice. Scale bar: 50 μm (top), 5 μm (bottom). (g) Western blotting with indicated antibodies of the spinal cord lysates from mice with indicated genotypes. (h) Expression of microglial markers IBA1 and CD68 in the spinal cords of mice with indicated genotypes. Scale bar: 50 μm (left), 5 μm (right).

**Figure S5. IFNγ deficiency suppresses the reactive astrocyte pathway in Arf1-ablated mice.**

(a) Representative brain sections from WT and Arf1-ablated mice were analyzed by immunohistochemistry for IBA1 (microglia) and GFAP (astrocytes). (b) Western blot detection of Arf1, GFAP, IBA1, and GAPDH in lysates of cerebellums, brain stems, and spinal cords from mice with the indicated genotypes. (c–e) Relative gene expression of the pan-astrocytes (c), A1 astrocytes (d), and A2 astrocytes (e) was assayed in the spinal cords of control and Arf1-ablated mice. (f) Representative brain sections from mice with the indicated genotypes were analyzed by immunofluorescence staining for GFAP (red), C3 (green), and Hoechst (blue). Scale bar, 100 μm (upper panel), 10 μm (lower panel). Data are shown as mean ± SEM. $^{*}P < 0.05$, $^{**}P < 0.01$, $^{***}P < 0.001$ using two-tailed t-test.

**Figure S6. Arf1 ablation induces the reactive astrocyte pathway through the IFNγ–Stat1 pathway.**

(a) Schematic map of brain domains with activated M1 microglia and A1 astrocytes in Arf1-ablated mice. (b) Western blot of phosphorylated Stat1 in lysates of cerebellum, medulla, and spinal cord from mice with the indicated genotypes. (c–g) IFNγ deficiency almost completely suppressed the neurodegenerative phenotypes of Arf1-ablated mice, including induction of IFNγ (c), TNFα (d), IL-1α (e), and C3 (f). C1q expression is not affected by Arf1 and IFNγ deficiencies (g). n = 5 per genotype, representing one of three independent experiments. Data are represented as mean ± SEM. *P < 0.05, **P < 0.01, ***P < 0.001 using two-tailed t-test.

**Figure S7. IFNγ deficiency did not affect Arf1-ablation-induced lipid peroxidation.**

(a) Electron microscopy sections of mouse spinal cords with the indicated genotypes. (b) Quantification of axon aggregates in the spinal cords of mice with the indicated genotypes (n = 20 images per genotype). (c) Quantification of lipid peroxidation based on malondialdehyde (MDA) levels in the spinal cord lysates of mice with the indicated genotypes (n=3 per group). (d) Immunofluorescence staining for BD-C11 to detect peroxided lipids in the white matter of spinal cords of mice with the indicated genotypes. Scale bar: 50 μm (top), 20 μm (bottom). (e) Quantification of lipid peroxidation based on BD-C11 levels in the spinal cords of mice with the indicated genotypes. Data are analyzed by two-tailed t-test and presented as mean ± SEM. *P < 0.05, **P < 0.01, ***P < 0.001.

**Figure S8. Treatment with the microglia inhibitor minocycline ameliorates the neurodegenerative phenotype caused by Arf1 ablation.**

(a–c) Neurological score (a), balance beam test (b), and body weight (c) of control and Arf1-ablated mice treated with saline or minocycline (n = 5 per genotype). (d) Representative brain sections from WT and Arf1-ablated mice were analyzed by immunohistochemistry for IBA1 (microglia) after treatment with saline or minocycline. Scale bar, 200 μm. (e, f) Number of IBA1+ cells was counted in the different brain areas and spinal cords of mice with the indicated genotypes and treatments (n = 13 per group). (g) Malondialdehyde (MDA) levels in spinal cord lysates from mice with the indicated genotypes and treatments (n = 5 per group). (h) Deletion of TLR4 in the Arf1-ablated mice does not rescue the neurodegenerative phenotypes (n = 5 per genotype). Data are shown as mean ± SD. *P < 0.05, **P < 0.01, ***P < 0.001 using two-tailed t-test.

**Figure S9. Arf1 deletion with *UBC-CreER* and *Nes-Cre* resulted in similar neurodegenerative phenotypes.**

Comparison of control (*Nes-Cre/Arf1*$^{f/+}$) and Arf1-deleted (*Nes-Cre/Arf1*$^{f/f}$) mice. (a–d) Elevated reactive A1 astrocytes (GFAP$^{+}$C3$^{+}$) (a and b) and M1 microglia (IBA1$^{+}$) (c and d) in the spinal cords of neuronal Arf1-deleted mice. Data are shown as mean ± SD. ***P < 0.001 using two-tailed t-test.

**Figure S10. Arf1 deletion with *UBC-CreER* or *Nes-Cre* generated similar neurodegenerative phenotypes.**

Comparison of control (*Nes-Cre/Arf1*$^{f/+}$) and Arf1-deleted (*Nes-Cre/Arf1*$^{f/f}$) mice. (a–d) Relative gene expression of the pan-Astrocytes (a), A1 astrocytes (b), A2 astrocytes (c), and other genes (d) in the spinal cords of control and Arf1-deleted mice. (e) Western blot detection of Arf1, GFAP, IBA1, and GAPDH in lysates of cerebellum, brain stem, and spinal cord from mice with the indicated genotypes. (f) Western blot of phosphorylated Stat1 in lysates of cerebellum, medulla, and spinal cord from mice with the indicated genotypes (n = 4 each genotype). Data are shown as mean ± SEM. *P < 0.05, **P < 0.01, ***P < 0.001 using two-tailed t-test.

**Figure S11. Arf1 deletion with *UBC-CreER* or *Thy1-CreER* resulted in similar neurodegenerative phenotypes.**

(a) Immunofluorescence staining for Mog, Mbp, and Hoechst in the spinal cords of control (*Thy1-CreER/Arf1*$^{f/+}$*)* and adult neuronal Arf1-deleted (*Thy1-CreER/Arf1*$^{f/f}$) mice. Scale bar: 50 µm (left), 20 µm (right). (b) Immunoblot of Arf1 and myelin markers Mog, Mbp, Cnp, and Mpz; neuron marker NeuN; astrocyte marker GFAP; and microglia marker IBA1 in the spinal cords of control (*Thy1-CreER/Arf1*$^{f/+}$) and adult neuronal Arf1-deleted (*Thy1-CreER/Arf1*$^{f/f}$) mice. (c) Representative spinal cord sections from mice with the indicated genotypes were analyzed by immunofluorescence staining for synaptophysin (red), PSD95 (green), and Hoechst (blue). Scale bar: 50 µm (top), 20 µm (bottom). (d) Quantification of PSD95-positive and synaptophysin-positive synapses in the spinal cords of mice with the indicated genotypes (n = 12). (e) Immunofluorescence staining for BD-C11 to detect peroxided lipids in the white matter of spinal cords of mice with the indicated genotypes. Scale bar: 50 µm (top), 20 µm (bottom). (f) Quantification of BD-C11 level in the spinal cords of mice with the indicated genotypes (n = 12). (g) Representative spinal cord sections from mice with the indicated genotypes were analyzed by immunohistochemistry for IBA1 (microglia) and GFAP (astrocytes). (h) Quantification of IBA1$^{+}$ microglia in different brain areas of mice with the indicated genotypes (n = 8 in each group). (i) Quantification of GFAP$^{+}$ astrocytes in different brain areas and spinal cords of mice with the indicated genotypes ($n = 8$ in each group). Data are shown as mean ± SD. *$P < 0.05$, **$P < 0.01$, ***$P < 0.001$ using two-tailed t-test.

**Figure S12. Arf1 knockout does not affect T and B cells in brain parenchyma.**

(a) Representative flow cytometry plots showing frequencies of $CD11b^+F4/80^+CD45^+$ monocytes and $CD11b^+Ly6G/C^+CD45^+$ macrophages in cerebellum and spinal cord tissues of control and Arf1-ablated mice. (b) Bar graph showing frequencies of immune cells in the spinal cord and cerebellum of control and Arf1-ablated mice ($n = 5$ per genotype). (c–f) Immunofluorescence staining for CD3, Mbp, and Hoechst (c); CNP, calreticulin, and Hoechst (d); HMGB1, Mbp, and Hoechst (e); and CD8α, Mog, and Hoechst (f) in spinal cord sections of mice with the indicated genotypes. Scale bar: 50 μm (left), 20 μm (right). Data are shown as mean ± SEM. *$P < 0.05$, **$P < 0.01$, ***$P < 0.001$ using two-tailed t-test.

**Figure S13. Arf1 knockout only affects γδ T cells in meninges.**

(a) Meninges were dissected, and single-cell suspensions were immunostained with immune-cell-specific makers for sorting and counting by flow cytometry. No significant changes were found in the indicated cell types between control and Arf1-ablated mice ($n = 5$ per genotype). (b–h) *Rag1* deficiency significantly suppressed the neurodegenerative phenotypes of Arf1-ablated mice, including demyelination (b), induction of $IBA1^+IFN\gamma^+$ cells (c), synapse loss (d), $IBA1^+$ microglia (e, f), and $GFAP^+$ astrocytes (g, h). Data are represented as mean ± SEM. *$P < 0.05$, **$P < 0.01$, ***$P < 0.001$ using two-tailed t-test.

**Figure S14. Injection of antibodies of IFNγ, VLA-4, and γδ T-cell receptor partially suppressed the phenotypes of Arf1-ablated mice.**

(a) Immunofluorescence staining for BD-C11 and Hoechst to detect peroxided lipids in the white matter of spinal cords of mice with the indicated genotypes. Scale bar: 50 μm (top), 20 μm (bottom). (b) Quantification of peroxided lipids based on BD-C11 levels in the spinal cords of mice with the indicated genotypes. (c–e) Injection of antibodies of IFNγ, VLA-4, and γδ T-cell receptor significantly suppressed the phenotypes of Arf1-ablated mice, including induction of IBA1+ microglia (c, d) and GFAP+ astrocytes (c, e). $n = 8$ per genotype. Data are represented as mean ± SEM. *$P < 0.05$, **$P < 0.01$, ***$P < 0.001$ using two-tailed t-test

**Figure S15. Arf1-ablated neurons release peroxided lipids and ATP.**

(a) Oil Red O staining of spinal cords from WT and Arf1-ablated mice. Scale bar: 50 μm. (b) Quantification of Oil-Red-O-positive cells in the different brain regions and the spinal cords of control and Arf1-ablated mice (n = 8). We found that, in comparison with those in brains of wild-type mice, lipid droplets were dramatically increased in brains of Arf1-ablated mice, particularly in the spinal cord and hindbrain areas (including cerebellum, pons, medulla, midbrain) but not in the forebrain areas (including the cerebral cortex and hippocampus). (c) BD-C11 immunostaining of peroxided lipids in spinal cords of control and Arf1-ablated mice. Scale bar: 50 μm (top), 10 μm (bottom). (d) Quantification of peroxided lipids in the spinal cord sections of WT and Arf1-ablated mice (n = 15 per group). (e) ELISA of aconitase activity in the spinal cord lysates of WT and Arf1-ablated mice (n = 5 per group). (f) Western blot detection of Arf1 in N2A cells. Sh-Arf11 and 2 significantly knocked down Arf1 expression. (g) Arf1 knockdown in N2A cells induced ATP expression. ELISA of ATP from culture medium of N2A cells transfected with sh-Scram or sh-Arf1. (h) Experimental setup for coculture of N2A and EOC 20 microglia cells to stain lipids. (i) Immunofluorescence staining for IBA1, BD-C11, and Hoechst in cocultured N2A and EOC 20 cells. N2A cells were first treated with sh-Scram or sh-Arf1 and oleic acid for 24 hours, washed two times with medium, and then added to EOC 20 cells for another 48 hours. Scale bar: 10 μm. (j) Quantitation of BD-C11-positive dots in EOC 20 cells (n = 20 per group). (k) ELISA of IL-1β in the EOC 20 cells treated with malondialdehyde (MDA) or 4-HNE (n = 4 per group). Data are represented as mean ± SEM. **P < 0.01, ***P < 0.001 using two-tailed t-test.

**Figure S16. AD4 and oxATP significantly suppressed the phenotypes associated with Arf1 ablation.**

Control (*UBC-CreER/Arf1$^{f/+}$*, WT) and Arf1-ablated (*UBC-CreER/Arf1$^{f/f}$*, Arf1$^{-/-}$) mice were treated with PBS, AD4, and oxATP and assayed for various phenotypes, as follows: (a) neurological score and balance beam test for mice with the indicated genotypes and treatments (n = 5 per genotype); (b) ELISA of aconitase activity in the spinal cord lysates of mice with the indicated genotypes and treatments (n = 5 per genotype and treatment); (c) malondialdehyde (MDA) levels in spinal cord lysates from mice with the indicated genotypes and treatments (n = 5 per condition); (d) IL-1β in spinal cord lysates from mice with the indicated genotypes and treatments (n = 5 per condition); (e) immunofluorescence staining for BD-C11 and Hoechst in spinal cord sections in mice with the indicated genotypes and treatments (n = 5 per condition), scale bars: 50 μm (top), 20 μm (bottom); (f) quantification of BD-C11 dots in panel e; (g–j)

TNFα (g), IFNγ (h), IL-1α (i), and C3 (j) in spinal cord lysates of mice with the indicated genotypes and treatments (n = 5 per condition); (k) immunofluorescence staining for IBA1, IFNγ, and Hoechst in spinal cord sections in mice with the indicated genotypes and treatments (n = 5 per condition), scale bar: 50 μm (top), 10 μm (bottom); (l) quantification of $IBA1^+$ microglia in panel k. Data are represented as mean ± SD. *P < 0.05, **P < 0.01, ***P < 0.001 using two-tailed t-test.

**Figure S17. AD4 and oxATP significantly suppressed the phenotypes associated with Arf1 ablation.**

Control (*UBC-CreER/Arf1*$^{f/+}$, WT) and *Arf1*-ablated (*UBC-CreER/Arf1*$^{f/f}$, $Arf1^{-/-}$) mice were treated with PBS, AD4, and oxATP and assayed for various phenotypes, as follows: (a) immunofluorescence staining for IBA1, IL-1β, and Hoechst in spinal cord sections in mice with the indicated genotypes and treatments, scale bar: 50 μm (top), 10 μm (bottom); (b) quantification of $IBA1^+IL\text{-}1\beta^+$ microglia in panel a (n = 5 per condition); (c) immunohistochemistry analysis for GFAP (astrocyte) in representative spinal cord sections from mice with the indicated genotypes and treatments, scale bars: 100 μm (top), 50 μm (bottom); (d) quantification of $GFAP^+$ astrocytes in different brain areas in mice with the indicated genotypes and treatments (n = 8 per genotype); (e–g) qPCR assay of chemokines in N2A cells (neuron) transfected with sh-Scram or sh-Arf1 (e), EOC 20 (microglia) treated with a medium of N2A cells transfected with sh-Scram or sh-Arf1 (f), and C8-D1A cells (astrocytes) treated with culture medium from panel f (g) (n = 3 per group). Data are represented as mean ± SEM. **P < 0.01, ***P < 0.001 using two-tailed t-test.

**Figure S18. Arf1-ablation-induced neurodegeneration is through the NLRP3 inflammasome.**

Control (*UBC-CreER/Arf1*$^{f/+}$, WT) and Arf1-ablated (*UBC-CreER/Arf1*$^{f/f}$, $Arf1^{-/-}$) mice were assayed for various phenotypes, as indicated below. (a–c) Body weight (a), neurological score (b), and balance beam test (c) of mice with the indicated genotypes and treatments (n = 5 per genotype). (d, e) Meninges were dissected from mice with the indicated genotypes, and single-cell suspensions were immunostained. γδ T cells were gated on live, single, $CD3^+$, $TCR\gamma\delta^+$ events (d) and counted by flow cytometry (e) (n = 3 mice per group). (f) Body weight of mice with the indicated genotypes (n = 5 per genotype). (g) Representative spinal cord sections from mice with the indicated genotypes were analyzed by immunohistochemistry for IBA1 (microglia) and GFAP (astrocytes). Scale bars: 50 μm. (h, i) Quantification of $IBA1^+$ microglia (h) and $GFAP^+$ astrocytes (i) in different brain areas in mice with the indicated genotypes. (j) Western blotting of spinal cord lysates from mice with the indicated antibodies and genotypes. (k, l) TNFα (k) and

IL-1α (l) in spinal cord lysates of mice with the indicated genotypes (n = 5 per condition). Data are represented as mean ± SEM. **P < 0.01, ***P < 0.001 using two-tailed t-test.

**Figure S19. Identification of a unique microglia signature associated with Arf1-knockout mice.**

(a) Uniform Manifold Approximation and Projection map of different clusters from single-microglia cell sequencing results in control, *Arf1*$^{-/-}$, *IFNg*$^{-/-}$, and *Arf1*$^{-/-}$*IFNg*$^{-/-}$ mice. (b) Cell ratio from cluster 1 to 12 in the four genotypes of microglia. (c) Network analysis of cluster 5 vs. cluster 1 in the microglia from *Arf1*$^{-/-}$ mice. (d) Representative top eight high-expression genes in cluster 5 compared with other clusters. (e, f) Heatmap showing from cluster 1 to cluster 5 down- (e) and up- (f) regulated genes compared with disease associated microglia genes in ALS and DAM. (g, h) Heat map showing the upregulated (g) and downregulated (h) genes in the *Arf1*$^{-/-}$ group compared to the other three groups. (i–l) Expression plot comparing *Arf1*$^{-/-}$ vs. wild-type (WT) single-cell RNA sequence data with published RNA-seq data of microglia in aging (i), ALS (g), disease-associated microglia (DAM) (k), and multiple sclerosis (l).

**Video S1.** Representative video of two-month-old *UBC-CreER/Arf1*$^{f/+}$ mice (WT) after tamoxifen (TAM) injection for 22 days.

**Video S2.** Representative video of two-month-old *UBC-CreER/Arf1*$^{f/f}$ mice (*Arf1*$^{-/-}$) after tamoxifen (TAM) injection for 22 days.

**Video S3.** Representative video of *Nes-Cre/Arf1*$^{f/+}$ mice (WT) 18 days after birth.

**Video S4.** Representative video of *Nes-Cre/Arf1*$^{f/f}$ mice (*Arf1*$^{-/-}$) 18 days after birth.

**Video S5.** Representative video of two-month-old *Thy1-CreER/Arf1*$^{f/+}$ mice (WT) after TAM injection for 23 days.

**Video S6.** Representative video of two-month-old *Thy1-CreER/Arf1*$^{f/f}$ mice (*Arf1*$^{-/-}$) after TAM injection for 23 days.

**Video S7.** Representative video of two-month-old *UBC-CreER/Arf1*$^{f/f}$*/IFNg*$^{-/-}$ mice (*Arf1*$^{-/-}$*/IFNg*$^{-/-}$) after TAM injection for 22 days.

**Figure S1**

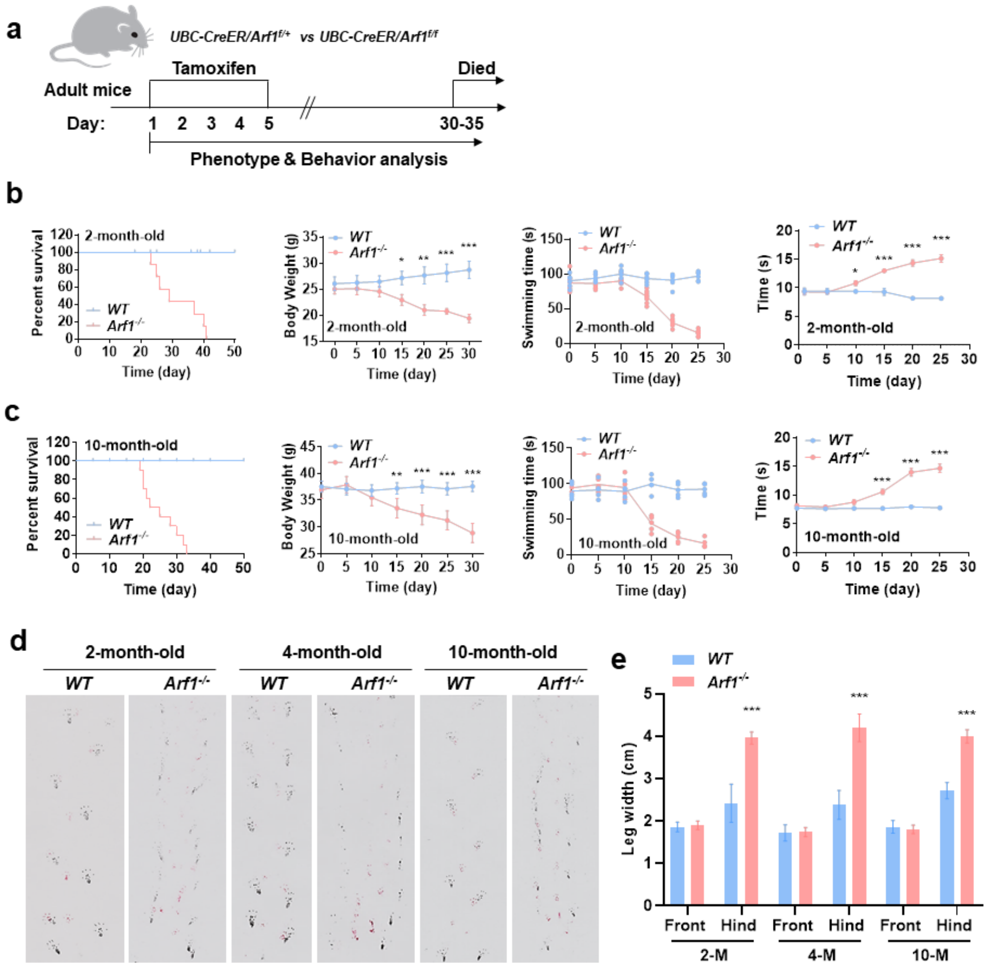
a
UBC-CreER/Arf1f/+ vs UBC-CreER/Arf1f/f
Tamoxifen
Died
Adult mice
Day: 1 2 3 4 5 30-35
Phenotype & Behavior analysis
b
2-month-old
Percent survival
Time (day)
WT
Arf1-/-
Body Weight (g)
Swimming time (s)
Time (s)
c
10-month-old
d
2-month-old
4-month-old
10-month-old
WT
Arf1-/-
e
Leg width (cm)
Front
Hind
2-M
4-M
10-M

### Figure S2

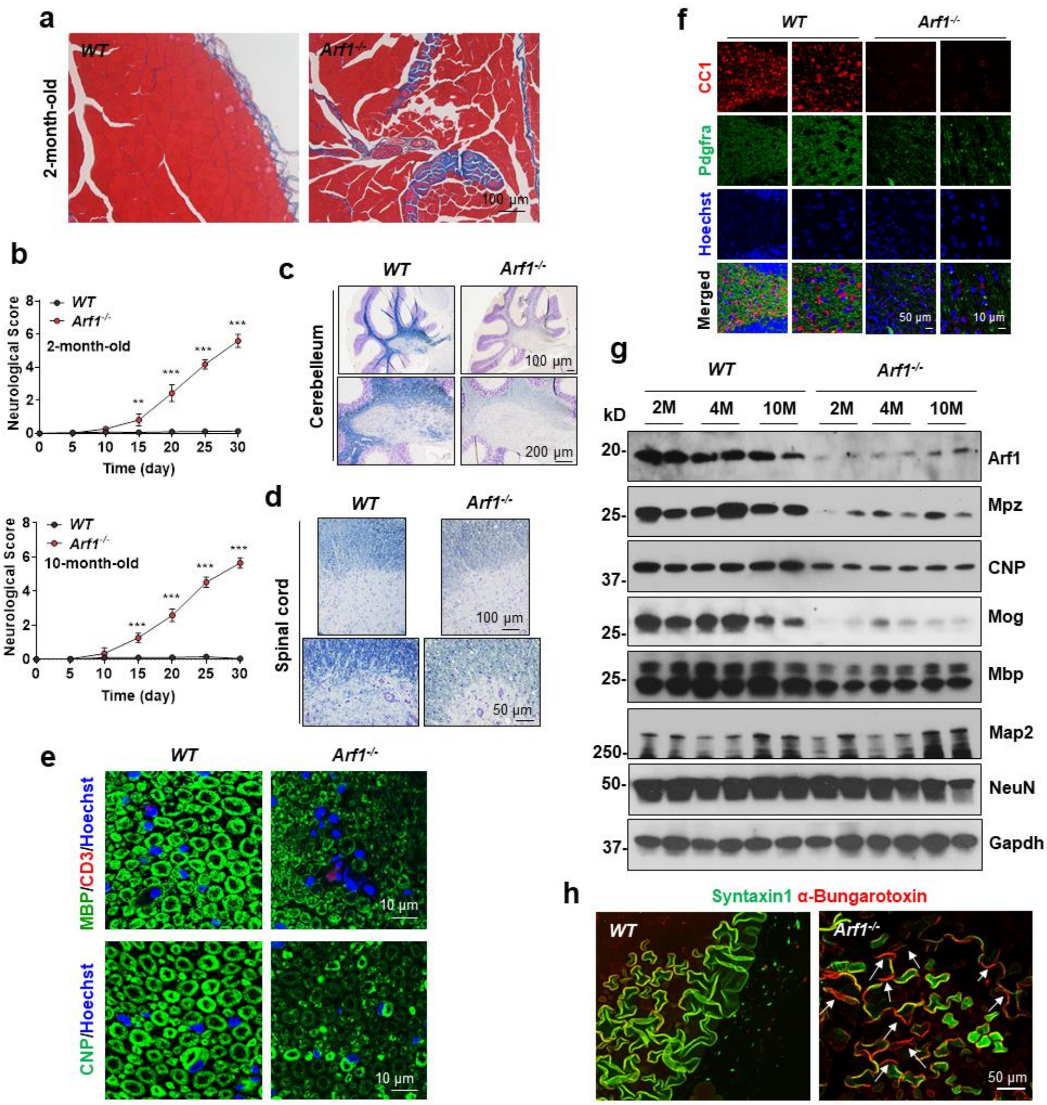

a
WT
Arf1-/-
2-month-old
100 µm
b
WT
Arf1-/-
2-month-old
Neurological Score
Time (day)
10-month-old
c
Cerebelleum
100 µm
200 µm
d
Spinal cord
50 µm
e
MBP/CD3/Hoechst
CNP/Hoechst
10 µm
f
CC1
Pdgfra
Hoechst
Merged
50 µm
g
kD
2M
4M
10M
Arf1
Mpz
CNP
Mog
Mbp
Map2
NeuN
Gapdh
h
Syntaxin1 α-Bungarotoxin
50 µm

#### Figure S3

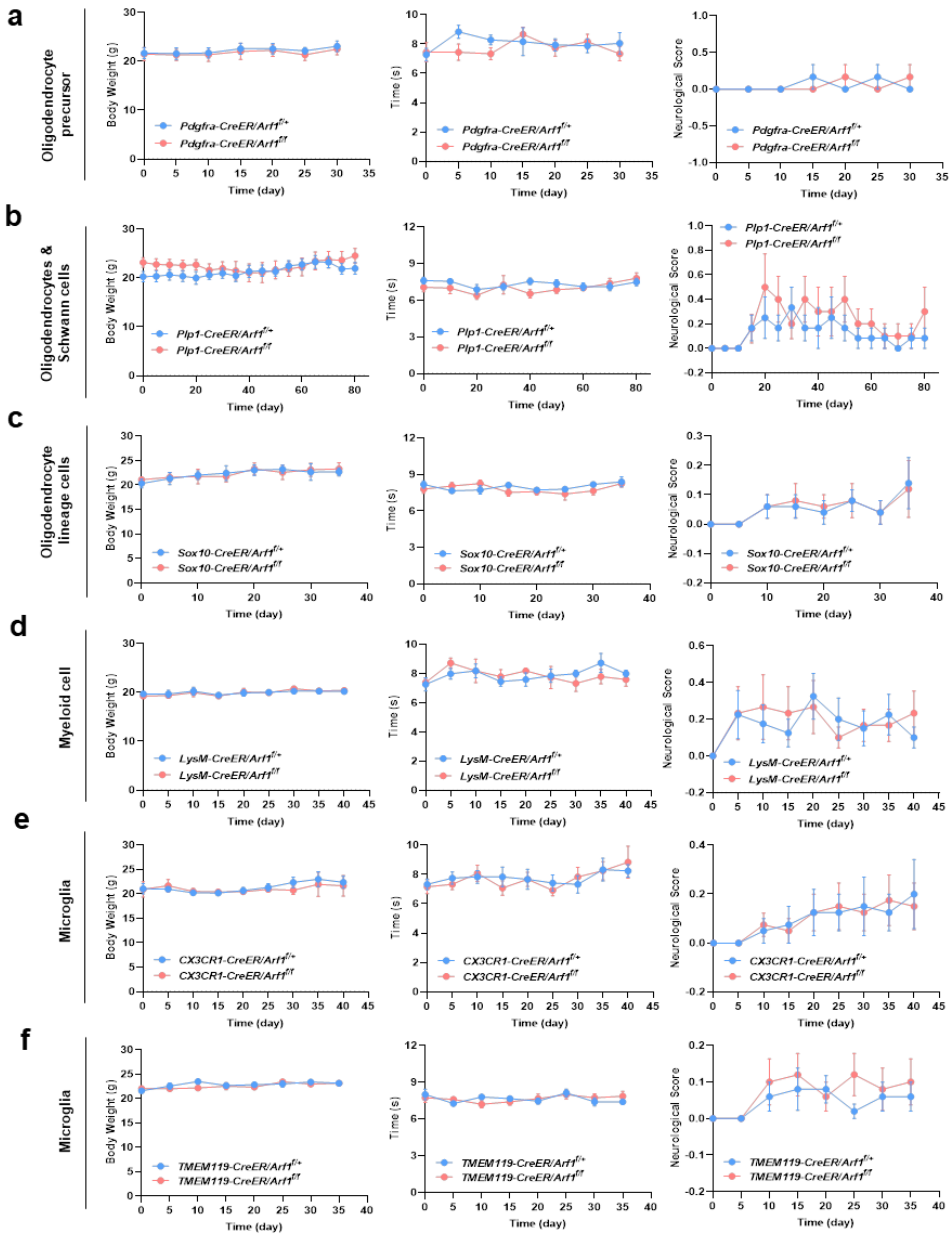

a
Oligodendrocyte precursor
Pdgfra-CreER/Arf1f/+
Pdgfra-CreER/Arf1f/f
b
Oligodendrocytes & Schwann cells
Plp1-CreER/Arf1f/+
Plp1-CreER/Arf1f/f
c
Oligodendrocyte lineage cells
Sox10-CreER/Arf1f/+
Sox10-CreER/Arf1f/f
d
Myeloid cell
LysM-CreER/Arf1f/+
LysM-CreER/Arf1f/f
e
Microglia
CX3CR1-CreER/Arf1f/+
CX3CR1-CreER/Arf1f/f
f
Microglia
TMEM119-CreER/Arf1f/+
TMEM119-CreER/Arf1f/f
Body Weight (g)
Time (s)
Neurological Score
Time (day)

### Figure S4

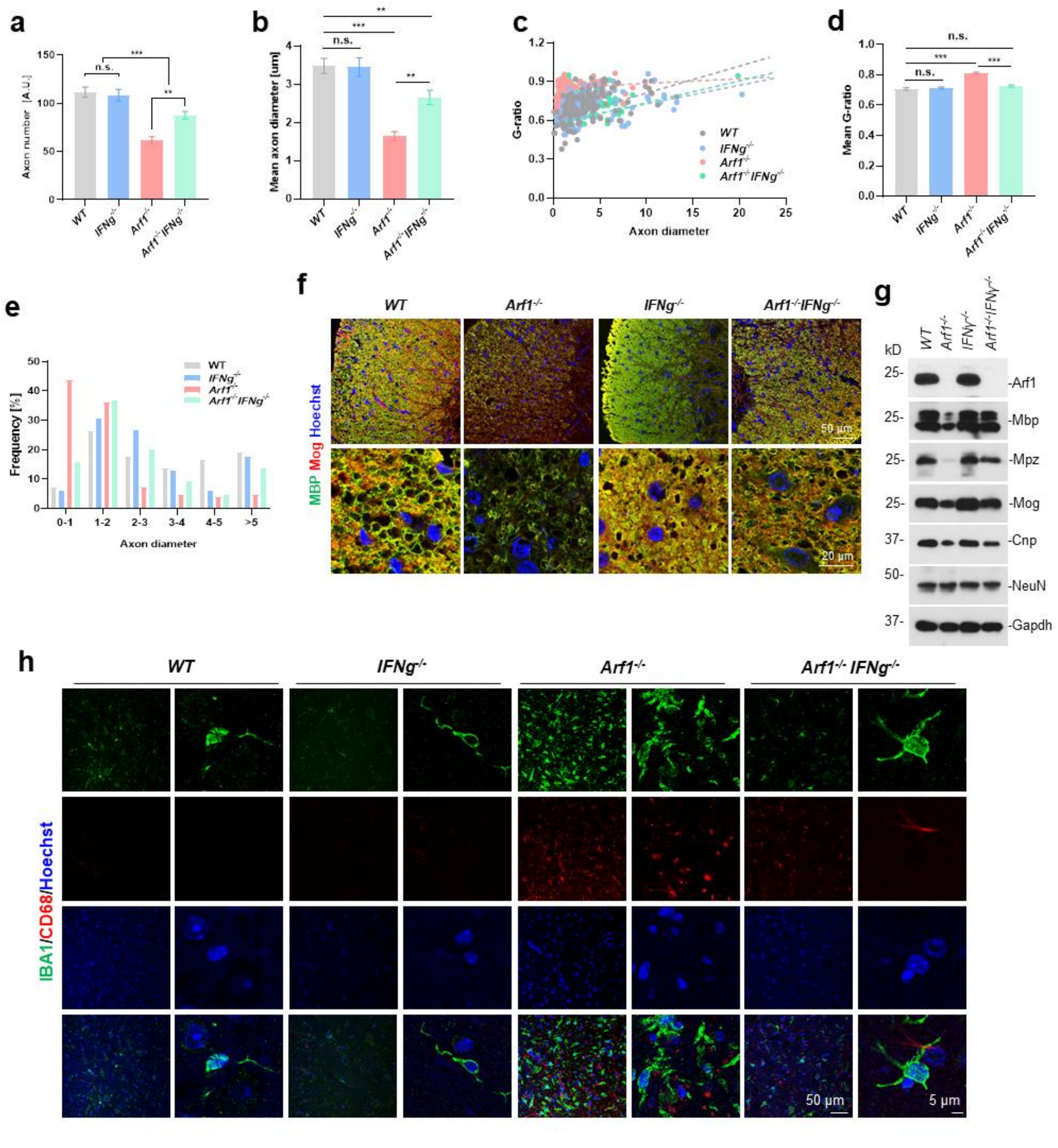

a
b
c
d
e
f
g
h
Axon number [A.U.]
Mean axon diameter [um]
G-ratio
Axon diameter
Mean G-ratio
Frequency [%]
WT
IFNg-/-
Arf1-/-
Arf1-/- IFNg-/-
n.s.
MBP Mog Hoechst
50 μm
20 μm
kD
-Arf1
-Mbp
-Mpz
-Mog
-Cnp
-NeuN
-Gapdh
IBA1/CD68/Hoechst
5 μm

**Figure S5**

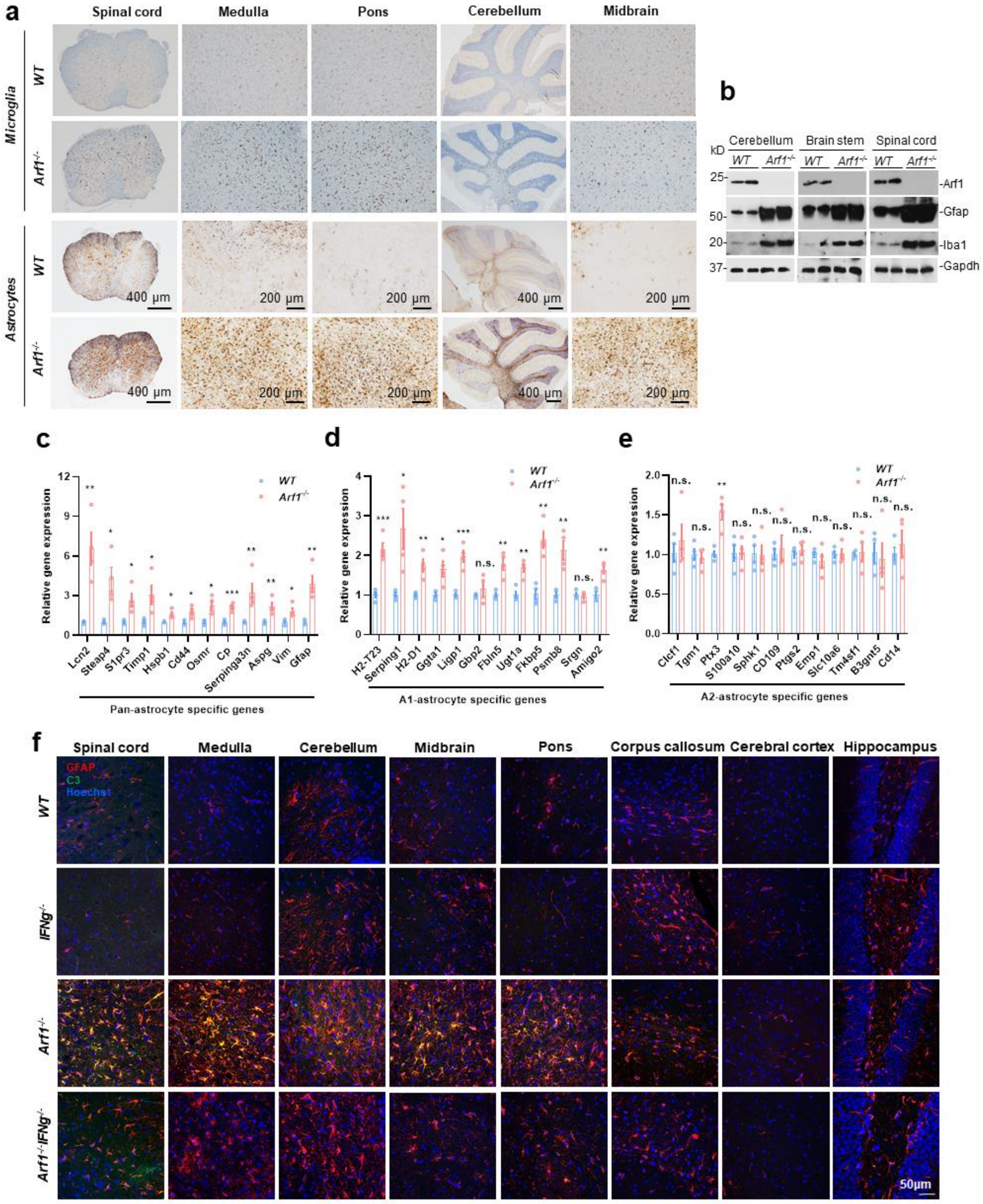
a
Spinal cord
Medulla
Pons
Cerebellum
Midbrain
Microglia
Astrocytes
WT
Arf1-/-
400 μm
200 μm
b
Cerebellum
Brain stem
Spinal cord
kD
25-
50-
20-
37-
-Arf1
-Gfap
-Iba1
-Gapdh
c
Relative gene expression
Pan-astrocyte specific genes
Lcn2
Steap4
S1pr3
Timp1
Hspb1
Cd44
Osmr
Cp
Serpinga3n
Aspg
Vim
Gfap
d
A1-astrocyte specific genes
H2-T23
Serping1
H2-D1
Ggta1
Ligp1
Gbp2
Fbln5
Ugt1a
Fkbp5
Psmb8
Srgn
Amigo2
e
A2-astrocyte specific genes
Clcf1
Tgm1
Ptx3
S100a10
Sphk1
CD109
Ptgs2
Emp1
Slc10a6
Tm4sf1
B3gnt5
Cd14
n.s.
f
Spinal cord
Medulla
Cerebellum
Midbrain
Pons
Corpus callosum
Cerebral cortex
Hippocampus
GFAP
C3
Hoechst
WT
IFNg-/-
Arf1-/-
Arf1-/-IFNg-/-
50μm

**Figure S6**

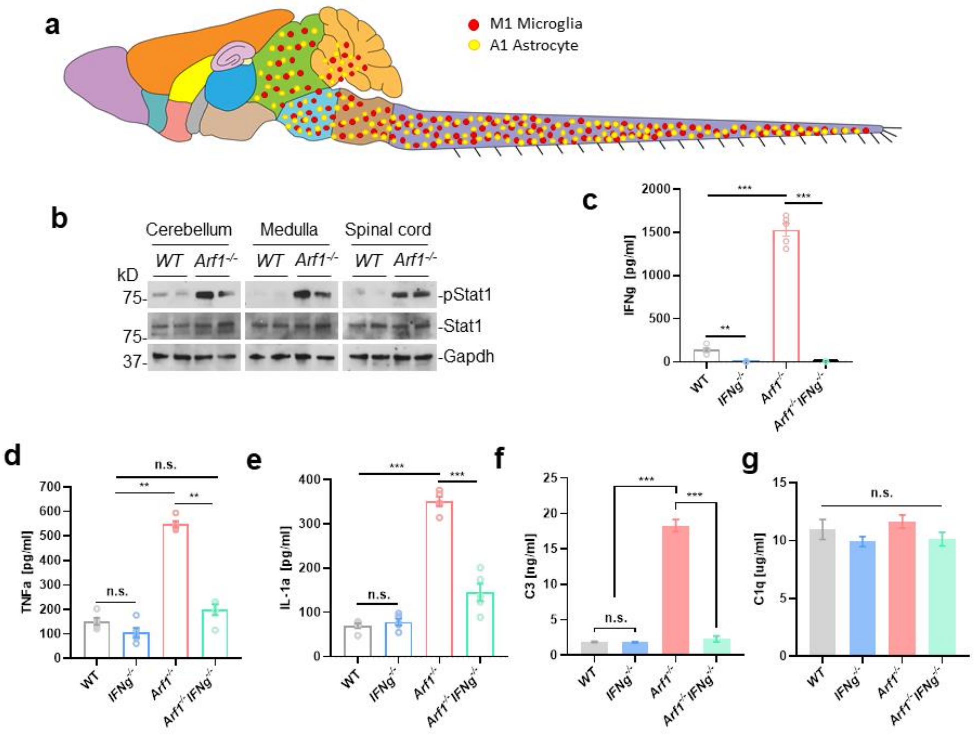

a
M1 Microglia
A1 Astrocyte
b
Cerebellum
Medulla
Spinal cord
WT
Arf1-/-
kD
75-
75-
37-
-pStat1
-Stat1
-Gapdh
c
IFNg [pg/ml]
d
TNFa [pg/ml]
e
IL-1a [pg/ml]
f
C3 [ng/ml]
g
C1q [ug/ml]
n.s.
WT
IFNg-/-
Arf1-/-
Arf1-/- IFNg-/-

#### Figure S7

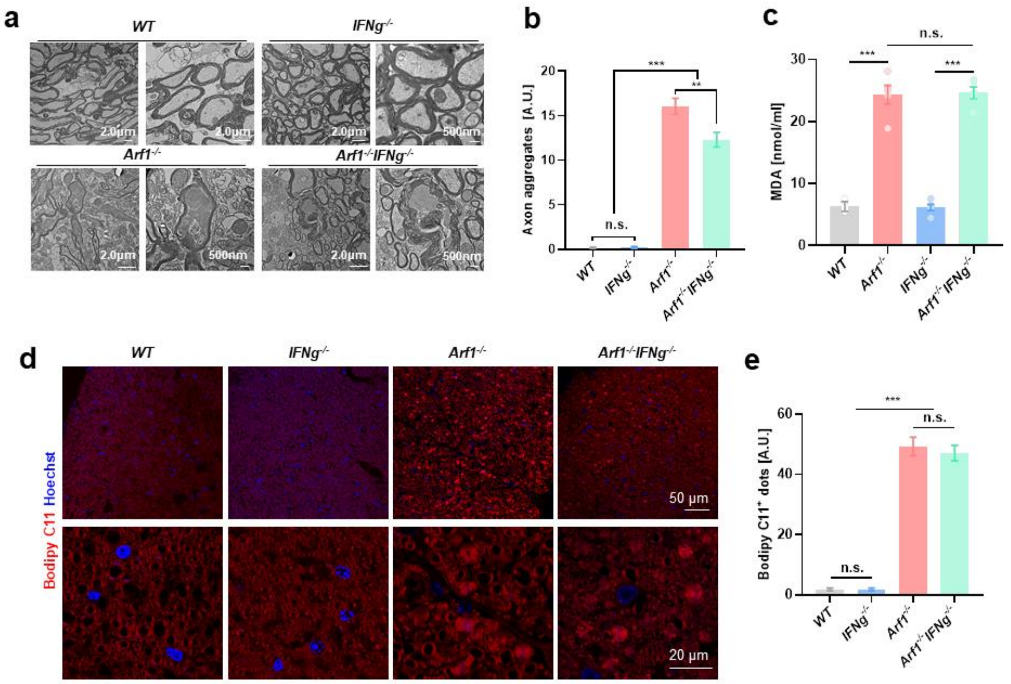
a
WT
IFNg-/-
Arf1-/-
Arf1-/-IFNg-/-
2.0μm
500nm
b
Axon aggregates [A.U.]
n.s.
***
**
WT
IFNg-/-
Arf1-/-
Arf1-/- IFNg-/-
c
MDA [nmol/ml]
n.s.
***
***
d
Bodipy C11 Hoechst
50 μm
20 μm
e
Bodipy C11+ dots [A.U.]

#### Figure S8

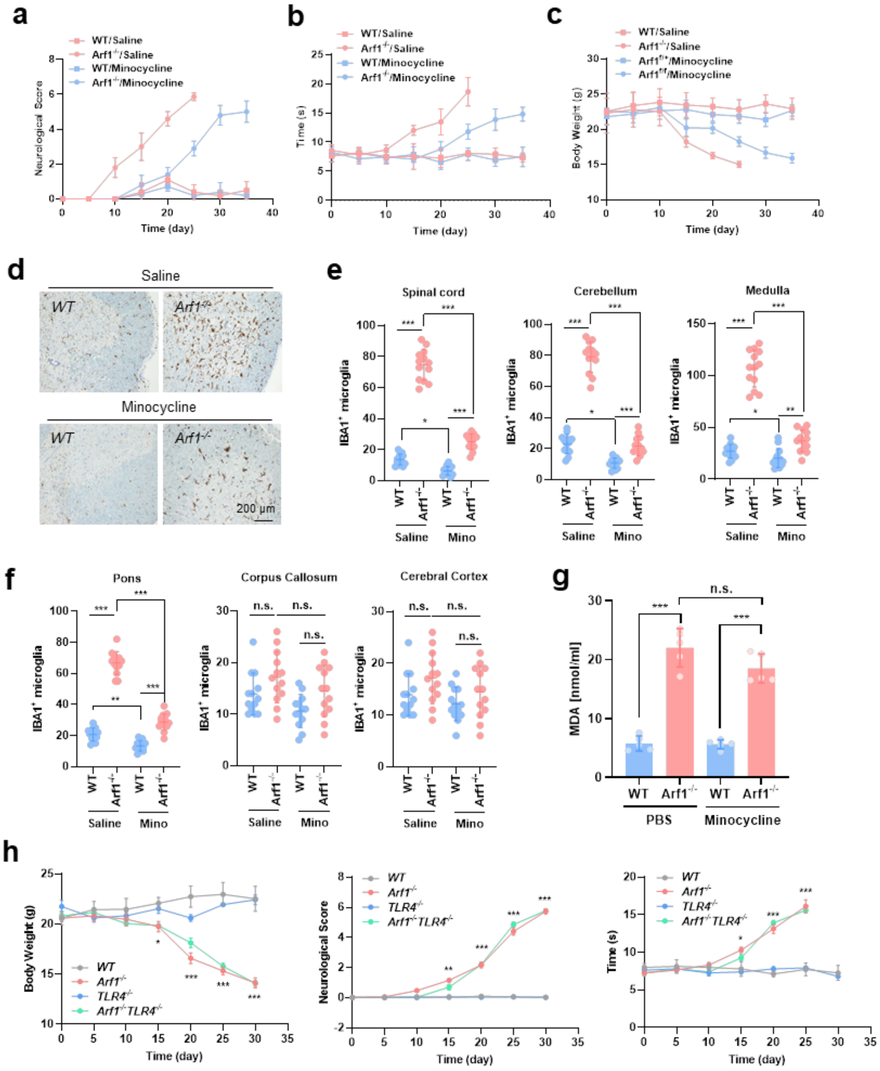
a
WT/Saline
Arf1-/-/Saline
WT/Minocycline
Arf1-/-/Minocycline
Neurological Score
Time (day)
b
Time (s)
c
Body Weight (g)
d
Saline
Minocycline
WT
Arf1-/-
200 μm
e
Spinal cord
Cerebellum
Medulla
IBA1+ microglia
Saline
Mino
f
Pons
Corpus Callosum
Cerebral Cortex
n.s.
g
MDA [nmol/ml]
PBS
Minocycline
h
WT
Arf1-/-
TLR4-/-
Arf1-/-TLR4-/-
Body Weight (g)
Neurological Score
Time (s)
Time (day)

### Figure S9

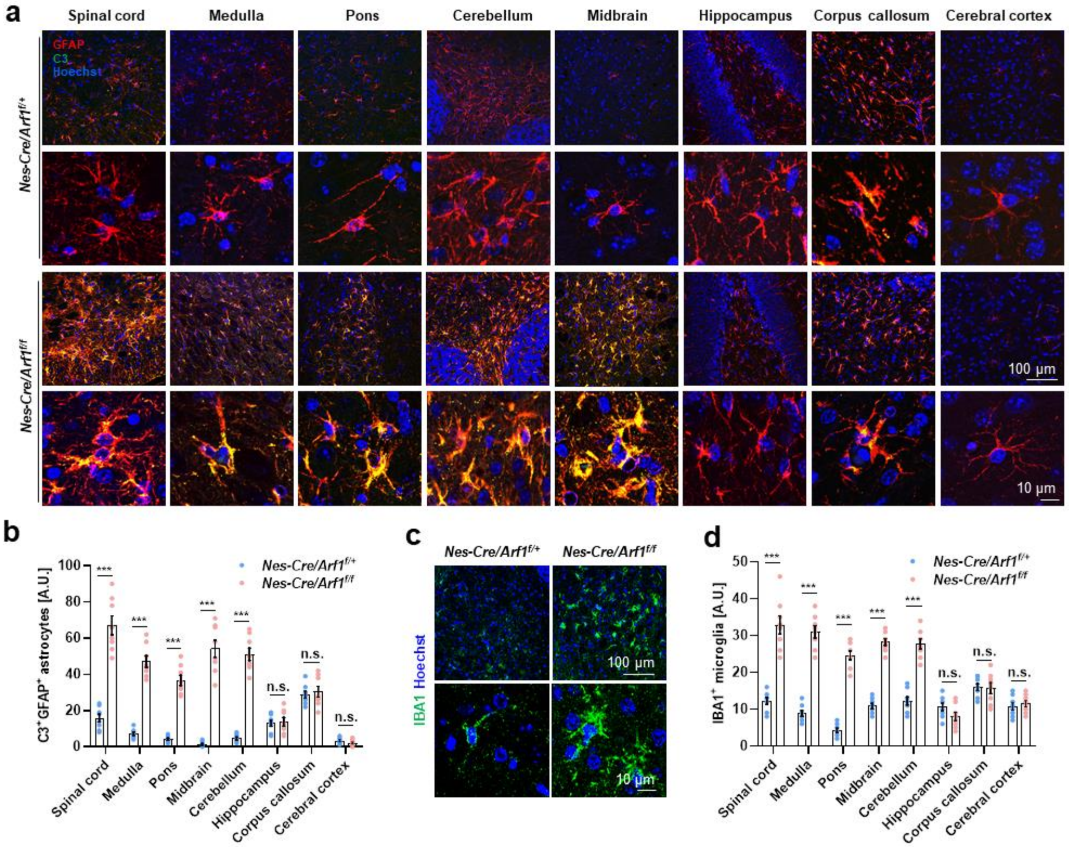
a
Spinal cord
Medulla
Pons
Cerebellum
Midbrain
Hippocampus
Corpus callosum
Cerebral cortex
GFAP
C3
Hoechst
Nes-Cre/Arf1f/+
Nes-Cre/Arf1f/f
100 μm
10 μm
b
C3+GFAP+ astrocytes [A.U.]
Nes-Cre/Arf1f/+
Nes-Cre/Arf1f/f
***
n.s.
c
Nes-Cre/Arf1f/+
Nes-Cre/Arf1f/f
IBA1 Hoechst
100 μm
10 μm
d
IBA1+ microglia [A.U.]
Nes-Cre/Arf1f/+
Nes-Cre/Arf1f/f
***
n.s.

**Figure S10**

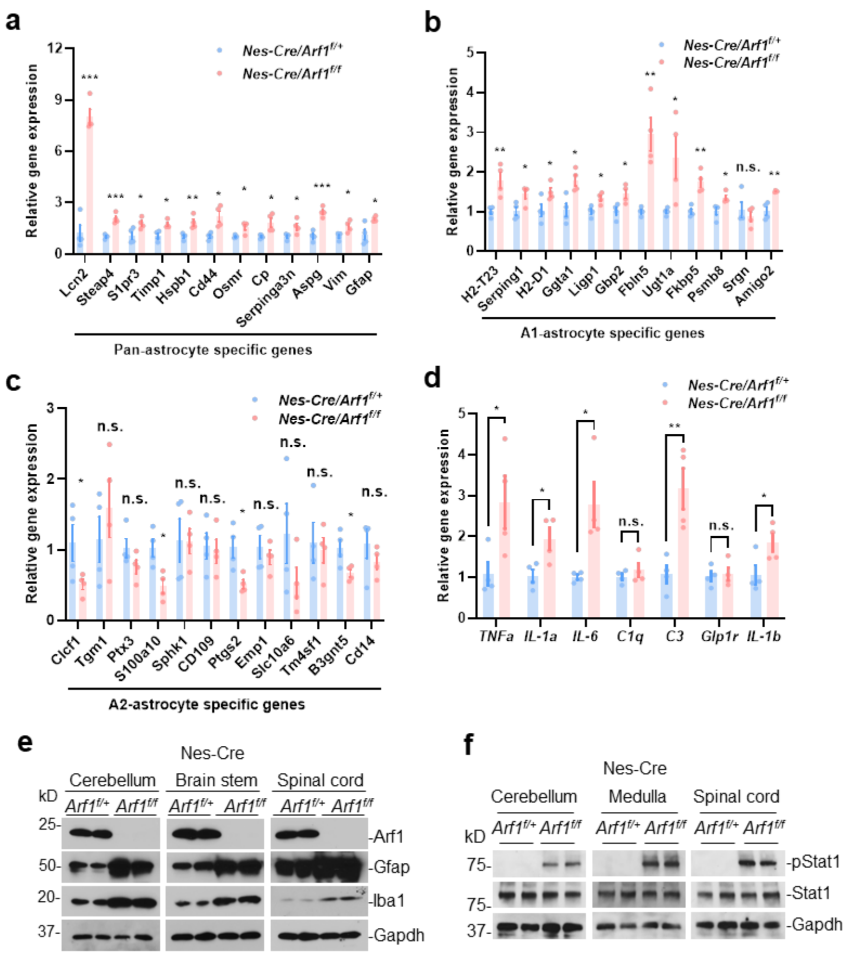
a
Relative gene expression
Nes-Cre/Arf1f/+
Nes-Cre/Arf1f/f
Lcn2
Steap4
S1pr3
Timp1
Hspb1
Cd44
Osmr
Cp
Serpinga3n
Aspg
Vim
Gfap
Pan-astrocyte specific genes
b
A1-astrocyte specific genes
H2-T23
Serping1
H2-D1
Ggta1
Ligp1
Gbp2
Fbln5
Ugt1a
Fkbp5
Psmb8
Srgn
Amigo2
c
A2-astrocyte specific genes
Clcf1
Tgm1
Ptx3
S100a10
Sphk1
CD109
Ptgs2
Emp1
Slc10a6
Tm4sf1
B3gnt5
Cd14
d
TNFa
IL-1a
IL-6
C1q
C3
Glp1r
IL-1b
e
Nes-Cre
Cerebellum
Brain stem
Spinal cord
kD
-Arf1
-Gfap
-Iba1
-Gapdh
f
Nes-Cre
Cerebellum
Medulla
Spinal cord
-pStat1
-Stat1
-Gapdh

#### Figure S11

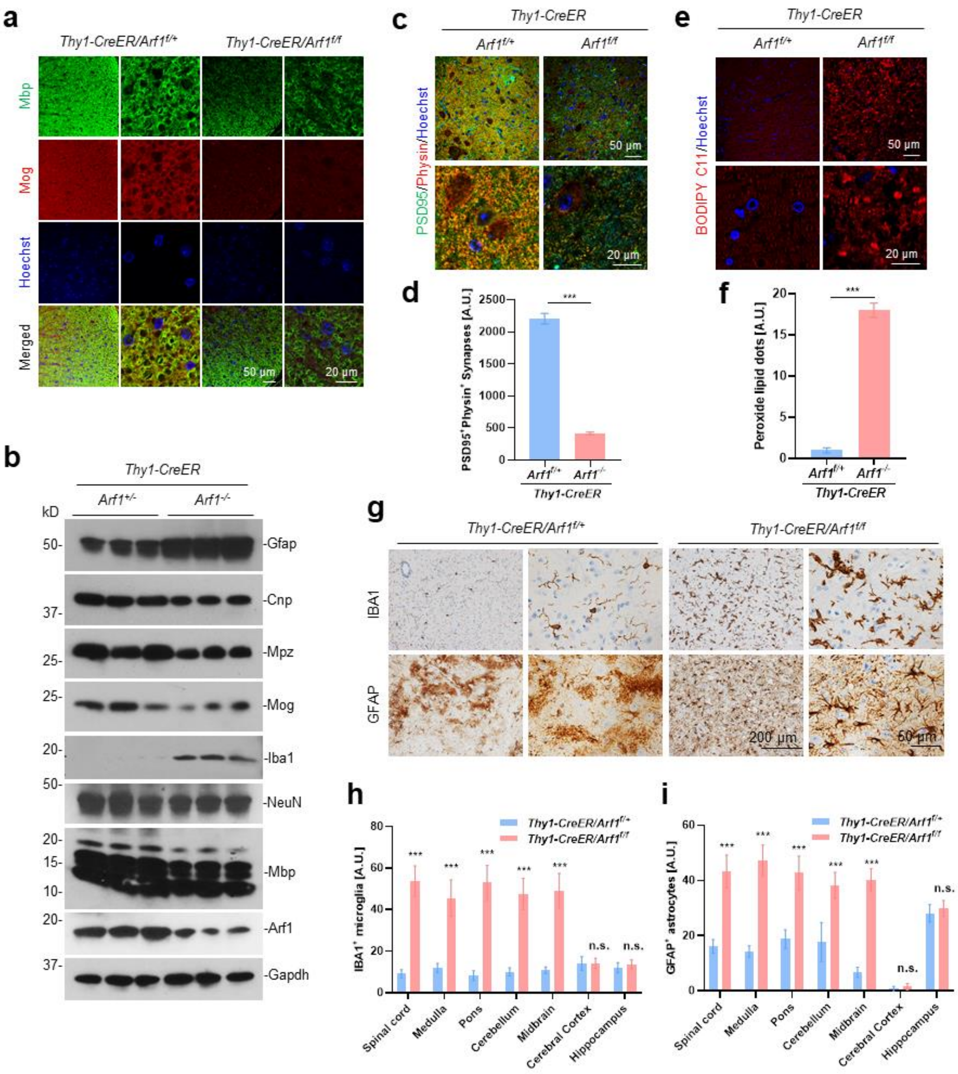
a
Thy1-CreER/Arf1f/+
Thy1-CreER/Arf1f/f
Mbp
Mog
Hoechst
Merged
50 μm
20 μm
b
Thy1-CreER
Arf1+/-
Arf1-/-
kD
-Gfap
-Cnp
-Mpz
-Mog
-Iba1
-NeuN
-Mbp
-Arf1
-Gapdh
c
Thy1-CreER
Arf1f/+
Arf1f/f
PSD95/Physin/Hoechst
d
PSD95+Physin+ Synapses [A.U.]
***
Arf1f/+
Arf1-/-
Thy1-CreER
e
Thy1-CreER
BODIPY C11/Hoechst
f
Peroxide lipid dots [A.U.]
g
Thy1-CreER/Arf1f/+
Thy1-CreER/Arf1f/f
IBA1
GFAP
200 μm
50 μm
h
IBA1+ microglia [A.U.]
i
GFAP+ astrocytes [A.U.]
n.s.
Spinal cord
Medulla
Pons
Cerebellum
Midbrain
Cerebral Cortex
Hippocampus

#### Figure S12

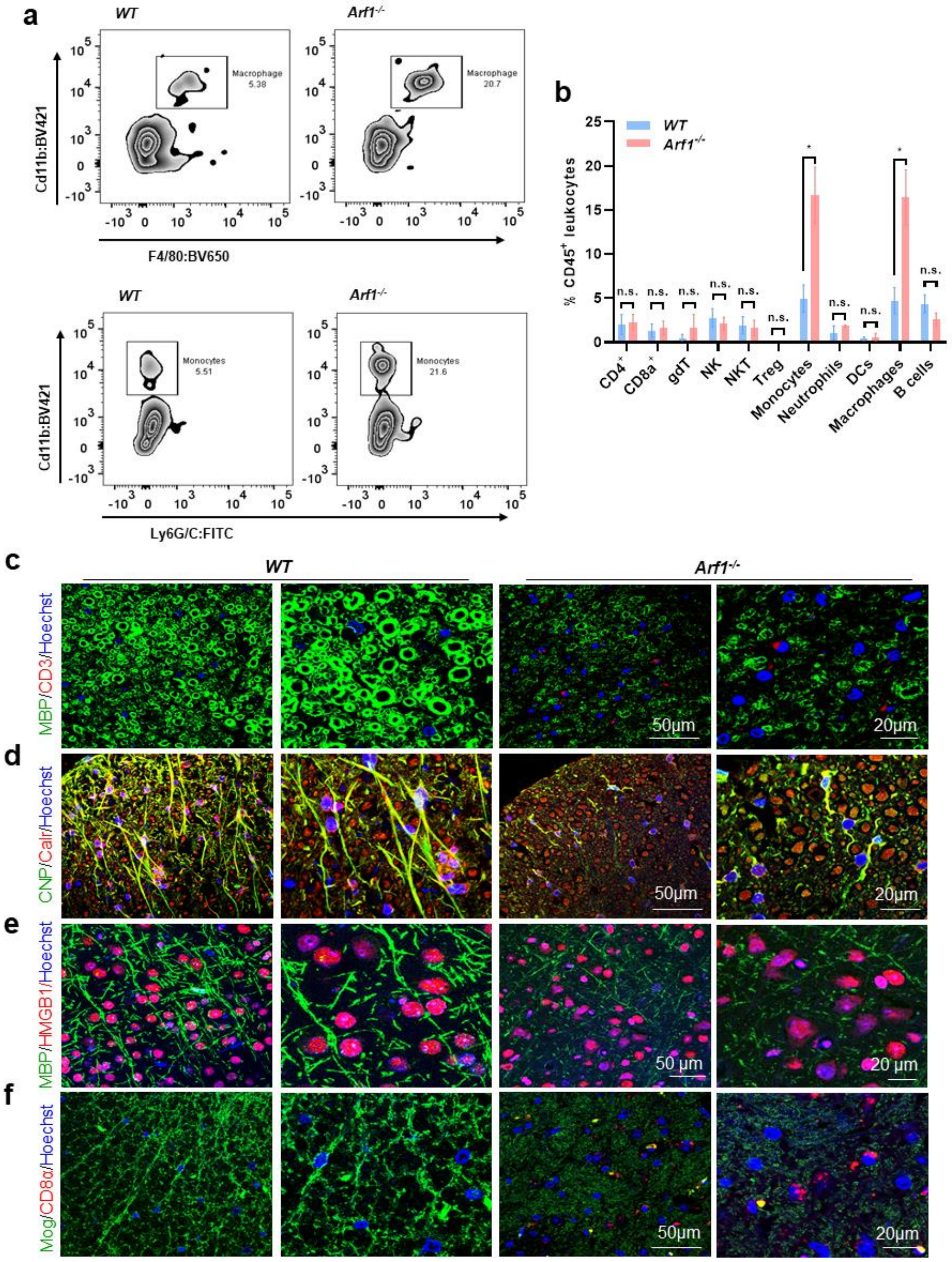

a
WT
Arf1-/-
Macrophage 5.38
Macrophage 20.7
Cd11b:BV421
F4/80:BV650
Monocytes 5.51
Monocytes 21.6
Ly6G/C:FITC
b
WT
Arf1-/-
% CD45+ leukocytes
n.s.
*
CD4+
CD8a+
gdT
NK
NKT
Treg
Monocytes
Neutrophils
DCs
Macrophages
B cells
c
MBP/CD3/Hoechst
d
CNP/Calr/Hoechst
e
MBP/HMGB1/Hoechst
f
Mog/CD8α/Hoechst
50µm
20µm

### Figure S13

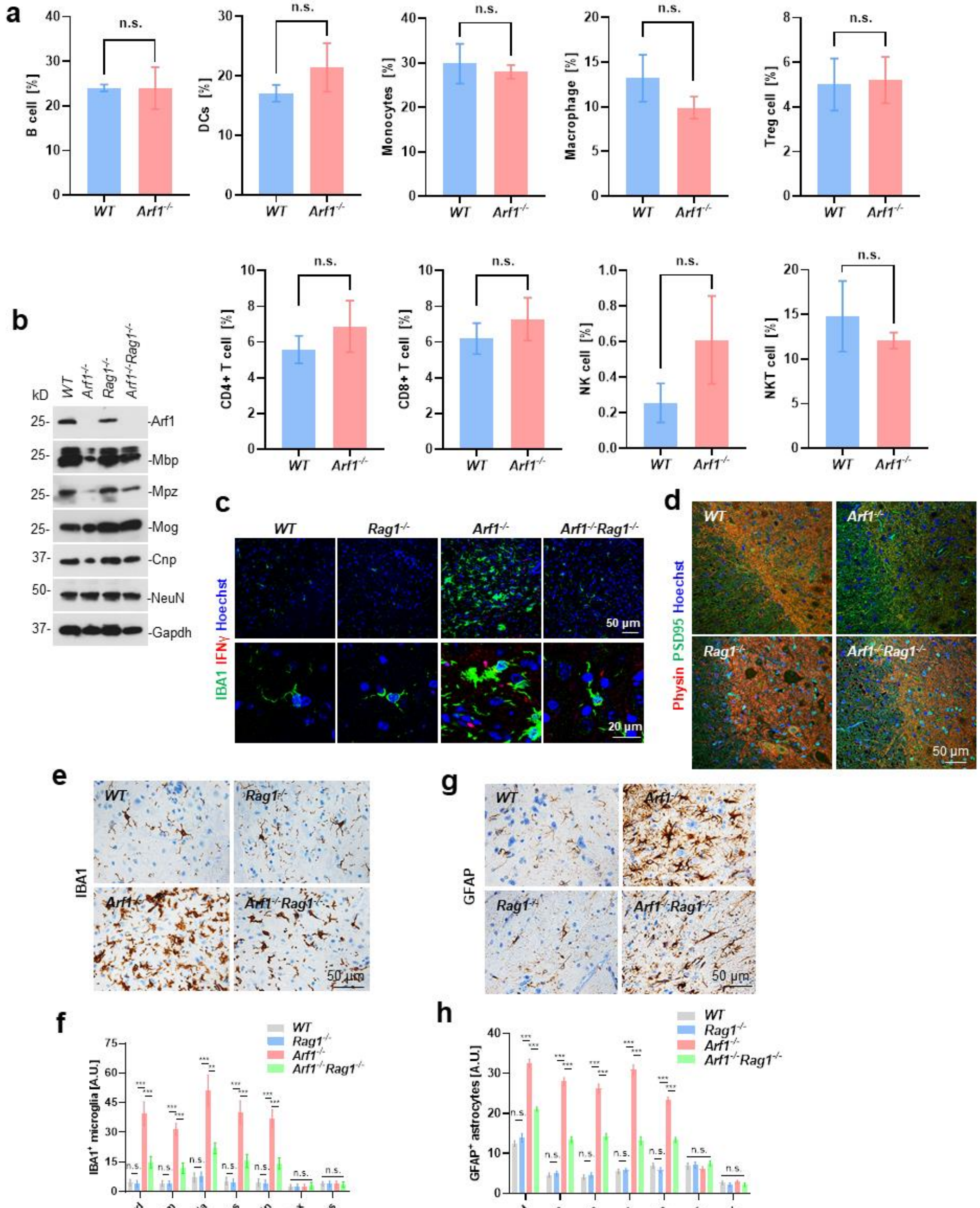

a
n.s.
B cell [%]
DCs [%]
Monocytes [%]
Macrophage [%]
Treg cell [%]
CD4+ T cell [%]
CD8+ T cell [%]
NK cell [%]
NKT cell [%]
WT
Arf1-/-
b
kD
WT
Arf1-/-
Rag1-/-
Arf1-/-Rag1-/-
-Arf1
-Mbp
-Mpz
-Mog
-Cnp
-NeuN
-Gapdh
c
IBA1 IFNγ Hoechst
50 µm
20 µm
d
Physin PSD95 Hoechst
50 µm
e
IBA1
g
GFAP
f
IBA1+ microglia [A.U.]
h
GFAP+ astrocytes [A.U.]
Spinal cord
Cerebellum
Medulla
Pons
Midbrain
Cerebral Cortex
Hippocampus

### Figure S14

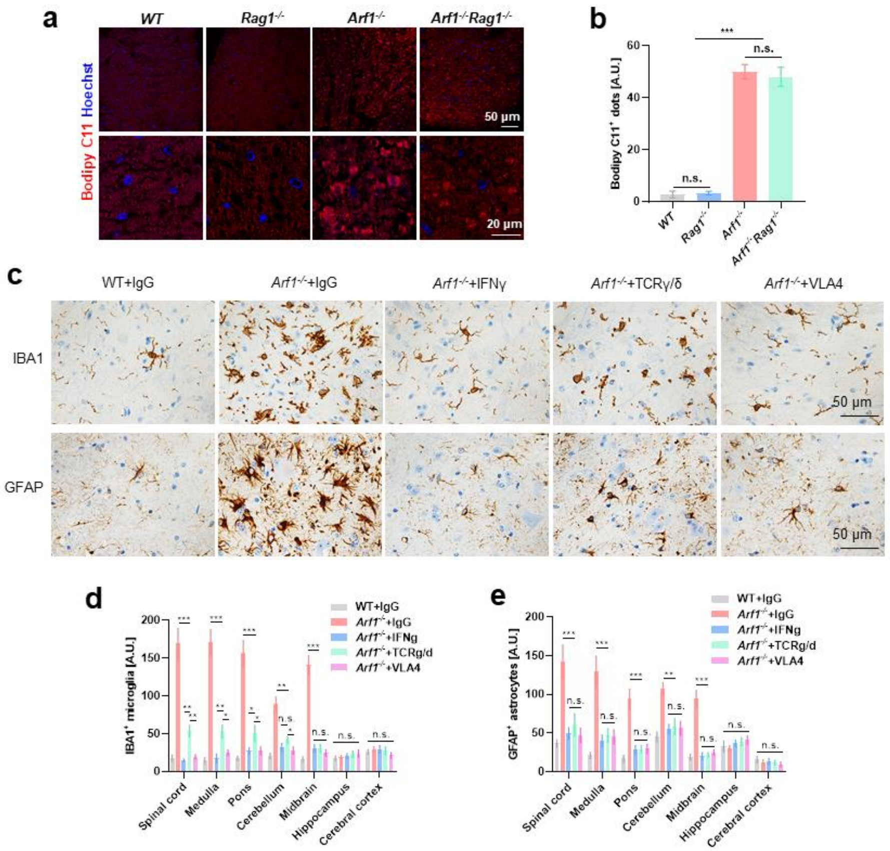
a
WT
Rag1-/-
Arf1-/-
Arf1-/-Rag1-/-
Bodipy C11 Hoechst
50 μm
20 μm
b
Bodipy C11+ dots [A.U.]
n.s.
***
c
WT+IgG
Arf1-/-+IgG
Arf1-/-+IFNγ
Arf1-/-+TCRγ/δ
Arf1-/-+VLA4
IBA1
GFAP
50 μm
d
IBA1+ microglia [A.U.]
WT+IgG
Arf1-/-+IgG
Arf1-/-+IFNg
Arf1-/-+TCRg/d
Arf1-/-+VLA4
Spinal cord
Medulla
Pons
Cerebellum
Midbrain
Hippocampus
Cerebral cortex
e
GFAP+ astrocytes [A.U.]

**Figure S15**

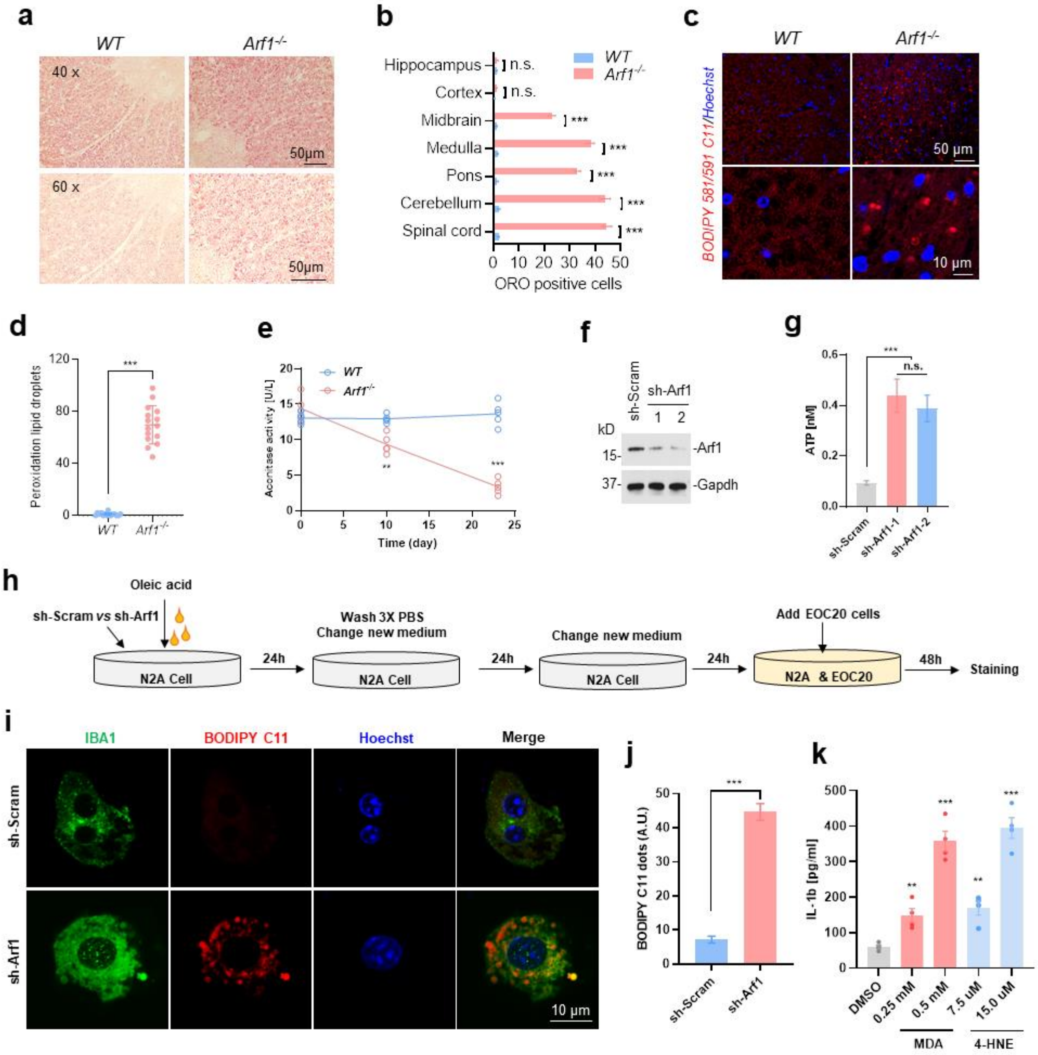

a
WT
Arf1-/-
40 x
60 x
50μm
b
Hippocampus
Cortex
Midbrain
Medulla
Pons
Cerebellum
Spinal cord
n.s.
***
WT
Arf1-/-
0 10 20 30 40 50
ORO positive cells
c
BODIPY 581/591 C11/Hoechst
50 μm
10 μm
d
Peroxidation lipid droplets
e
Aconitase activity [U/L]
Time (day)
f
sh-Scram
sh-Arf1
1 2
kD
15-
37-
-Arf1
-Gapdh
g
ATP [nM]
n.s.
sh-Scram
sh-Arf1-1
sh-Arf1-2
h
Oleic acid
sh-Scram vs sh-Arf1
N2A Cell
24h
Wash 3X PBS
Change new medium
Change new medium
Add EOC20 cells
N2A & EOC20
48h
Staining
i
IBA1
BODIPY C11
Hoechst
Merge
sh-Scram
sh-Arf1
10 μm
j
BODIPY C11 dots (A.U.)
k
IL-1b [pg/ml]
DMSO
0.25 mM
0.5 mM
7.5 uM
15.0 uM
MDA
4-HNE

**Figure S16**

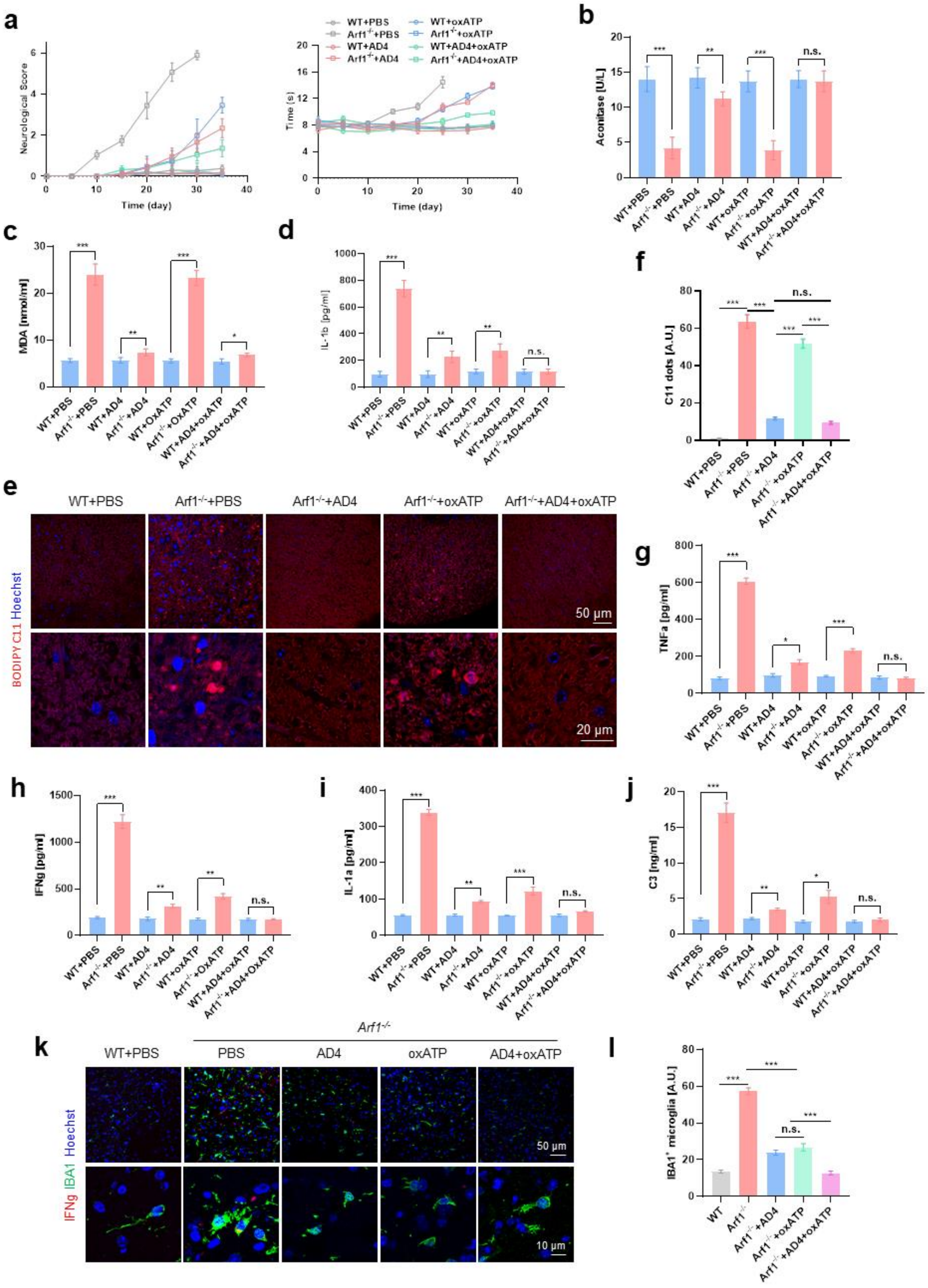
a
b
c
d
e
f
g
h
i
j
k
l
Neurological Score
Time (day)
Time (s)
WT+PBS
Arf1-/-+PBS
WT+AD4
Arf1-/-+AD4
WT+oxATP
Arf1-/-+oxATP
WT+AD4+oxATP
Arf1-/-+AD4+oxATP
Aconitase [U/L]
MDA [nmol/ml]
IL-1b [pg/ml]
C11 dots [A.U.]
BODIPY C11 Hoechst
50 μm
20 μm
TNFa [pg/ml]
IFNg [pg/ml]
IL-1a [pg/ml]
C3 [ng/ml]
Arf1-/-
PBS
AD4
oxATP
AD4+oxATP
IFNg IBA1 Hoechst
10 μm
IBA1+ microglia [A.U.]
n.s.

#### Figure S17

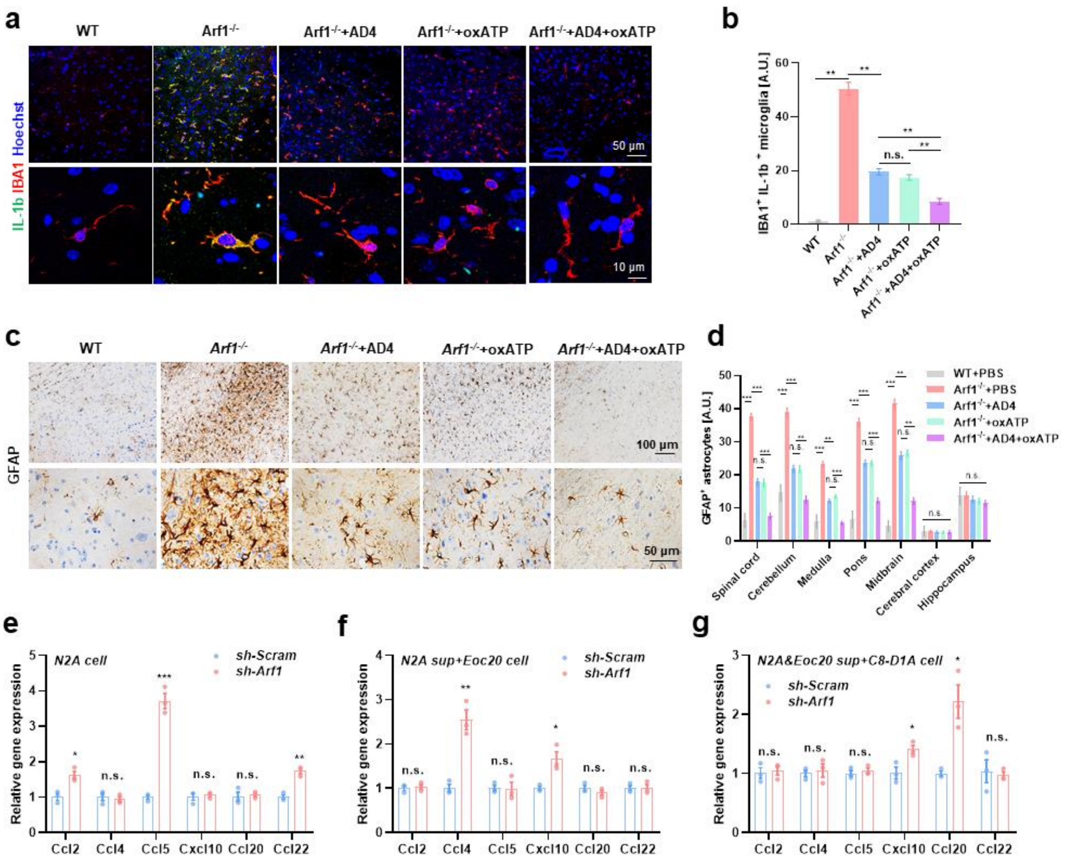
a
WT
Arf1-/-
Arf1-/-+AD4
Arf1-/-+oxATP
Arf1-/-+AD4+oxATP
IL-1b IBA1 Hoechst
50 μm
10 μm
b
IBA1+ IL-1b+ microglia [A.U.]
c
GFAP
100 μm
50 μm
d
GFAP+ astrocytes [A.U.]
WT+PBS
Arf1-/-+PBS
Arf1-/-+AD4
Arf1-/-+oxATP
Arf1-/-+AD4+oxATP
Spinal cord
Cerebellum
Medulla
Pons
Midbrain
Cerebral cortex
Hippocampus
e
N2A cell
sh-Scram
sh-Arf1
Relative gene expression
Ccl2
Ccl4
Ccl5
Cxcl10
Ccl20
Ccl22
f
N2A sup+Eoc20 cell
g
N2A&Eoc20 sup+C8-D1A cell

### Figure S18

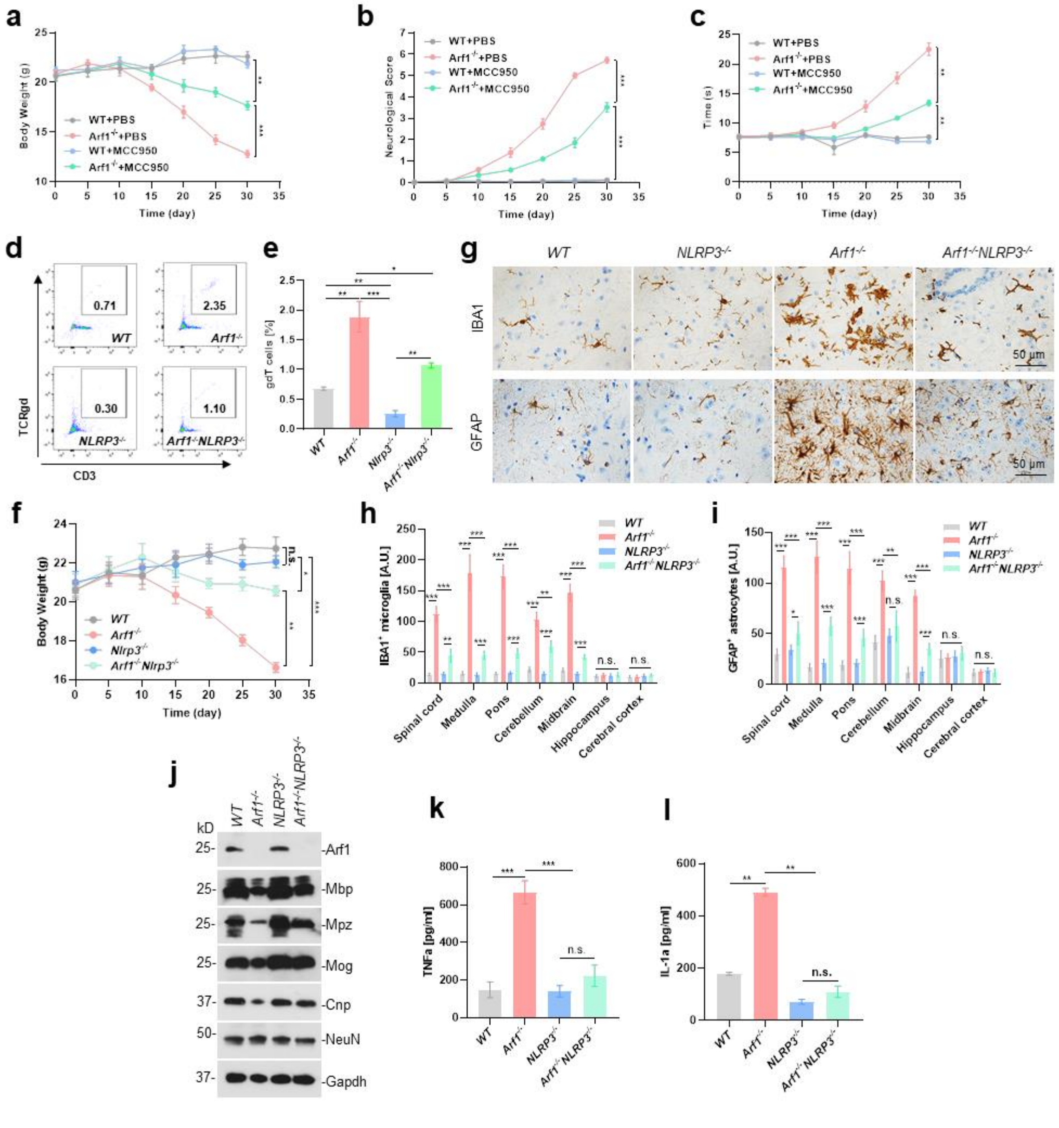
a
b
c
d
e
f
g
h
i
j
k
l
WT+PBS
Arf1-/-+PBS
WT+MCC950
Arf1-/-+MCC950
Body Weight (g)
Neurological Score
Time (s)
Time (day)
TCRgd
CD3
gdT cells [%]
WT
NLRP3-/-
Arf1-/-
Arf1-/-NLRP3-/-
IBA1
GFAP
50 μm
IBA1+ microglia [A.U.]
GFAP+ astrocytes [A.U.]
Spinal cord
Medulla
Pons
Cerebellum
Midbrain
Hippocampus
Cerebral cortex
-Arf1
-Mbp
-Mpz
-Mog
-Cnp
-NeuN
-Gapdh
TNFa [pg/ml]
IL-1a [pg/ml]

**Figure S19**

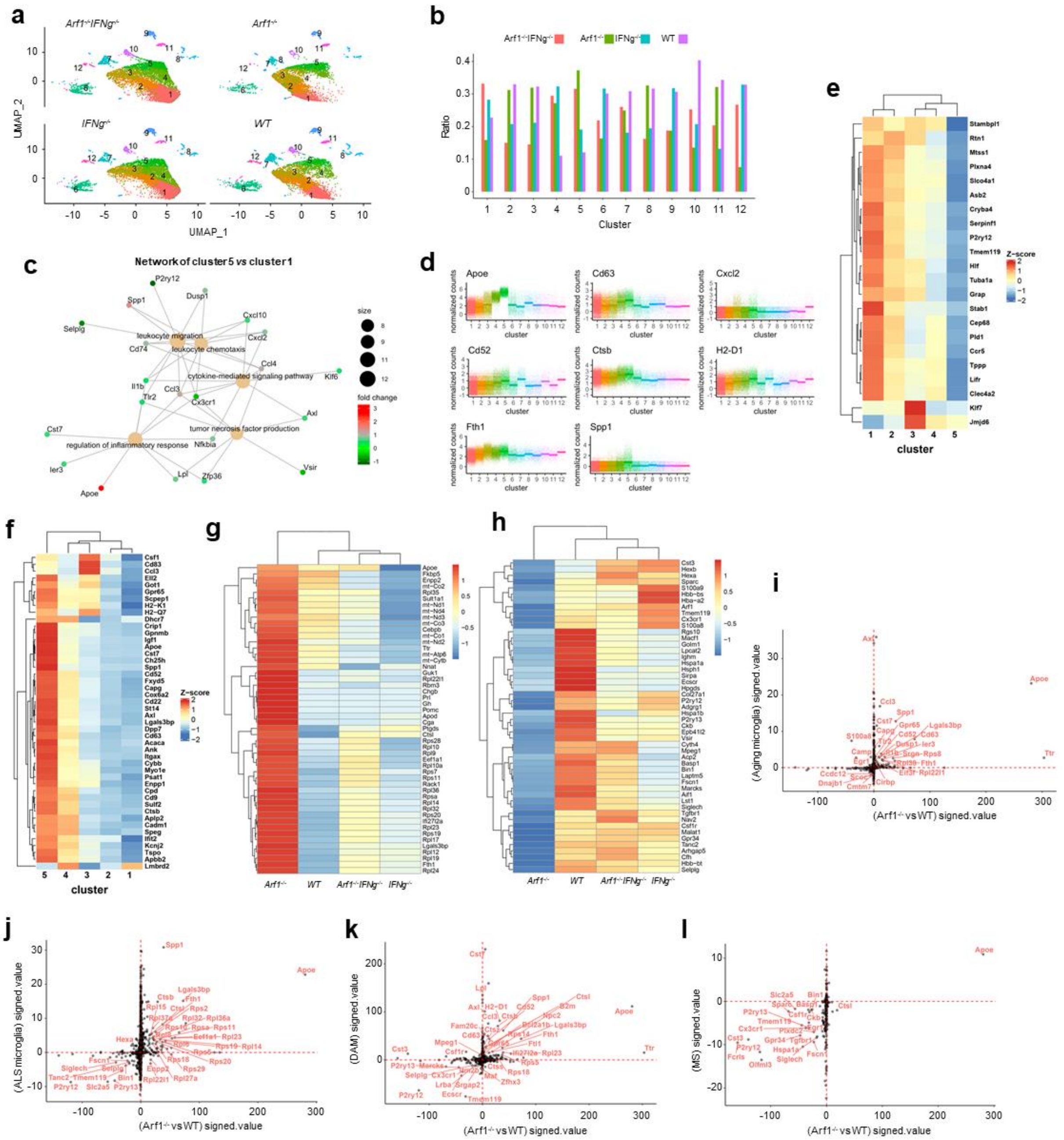

### Supplemental Table

**Supplemental Table 1. Primers used in this paper**

| Primer | Sequence |
|---|---|
| Ccl2-F | 5’-TCATGCTTCTGGGCCTGCTG-3’ |
| Ccl2-R | 5’-CTTTGGGACACCTGCTGCTG-3’ |
| Ccl3-F | 5’-CACTGCCCTTGCTGTTCTTC-3’ |
| Ccl3-R | 5’-CACCTGGCTGGGAGCAAAGG-3’ |
| Ccl4-F | 5’-GAAGCTCTGCGTGTCTGCCCT-3’ |
| Ccl4-R | 5’-CCACAGCTGGCTTGGAGCAA-3’ |
| Ccl5-F | 5’-CACCATCATCCTCACTGCAG-3’ |
| Ccl5-R | 5’-CTTCGAGTGACAAACACGAC-3’ |
| Cxcl9-F | 5’-GCATCATCTTCCTGGAGCA-3’ |
| Cxcl9-R | 5’-CTAGGCAGGTTTGATCTCCG-3’ |
| Cxcl10-F | 5’-TGAACCCAAGTGCTGCCGTC-3’ |
| Cxcl10-R | 5’-CATCGTGGCAATGATCTCAA-3’ |
| Cxcl11-F | 5’-GTCACAGCCATAGCCCTGGC-3’ |
| Cxcl11-R | 5’-CCTTCATAGTAACAATCACT-3’ |
| Ccl17-F | 5’-CTTCACCTCAGCTTTTGGTA-3’ |
| Ccl17-R | 5’-TACCAGCTCACCAACTTCCTG-3’ |
| Ccl20-F | 5’-GTCTGCTCTTCCTTGCTTTG-3’ |
| Ccl20-R | 5’-TGATAGCATTAATGTCACAAG-3’ |
| Ccl22-F | 5’-GGCTACCCTGCGTGTCCCACT-3’ |
| Ccl22-R | 5’-GGTCCAGAAGAACTCCTTCA-3’ |
| Ccl28-F | 5’-GCAGCAAGCAGGGCTCACA-3’ |
| Ccl28-R | 5’-GGATGACAGCAGCCAGGTCG-3’ |
| TNFα-F | 5’-CTACACAGAAGTTCCCAAAT-3’ |
| TNFα-R | 5’-CCCTTTCCTCCCAAACCAA-3’ |
| IL-1α-F | 5’-GCACCTTACACCTACCAGAGT-3’ |
| IL-1α-R | 5’-AAACTTCTGCCTGACGAGCTT-3’ |
| IL6-F | 5’-TAGTCCTTCCTACCCCAATTTCC-3’ |
| IL6-R | 5’-TTGGTCCTTAGCCACTCCTTC-3’ |
| IL-1β-F | 5’-GCTGCTTCCAAACCTTTGACC-3’ |
| IL-1β-R | 5’-GGTGCTCATGTCCTCATCCTGG-3’ |
| C1q-F | 5’-TCTGCACTGTACCCGGCTA-3’ |
| C1qa-R | 5’-CCCTGGTAAATGTGACCCTTTT-3’ |
| C3-F | 5’-CCAGCTCCCCATTAGCTCTG-3’ |
| C3-R | 5’-GCACTTGCCTCTTTAGGAAGTC-3’ |
| Glp1r-F | 5’-ACGGTGTCCCTCTCAGAGAC-3’ |
| Glp1r-R | 5’-ATCAAAGGTCCGGTTGCAGAA-3’ |
| Gapdh-F | 5’-GTCCCAGCTTAGGTTCATCA-3’ |
| Gapdh-R | 5’-GAGGTCAATGAAGGGGTCGT-3’ |

#### Supplemental Table 2. Information for Human Specimens

| SID | GUID | BW | DISORDER | AGE YEARS | AGE DAYS | SEX | RACE | HIV | HBsAG | PMINTERVAL |
|---|---|---|---|---|---|---|---|---|---|---|
| 1491 | NDAR_INVBP614GMR | N/A | MS | 55 | 14 | Male | White | N/A | N/A | 3 |
| 5274 | NDAR_INVTZ491ZPW | N/A | Normal | 64 | 187 | Female | White | Negative | Negative | 20 |
| 5487 | NDAR_INVZZ586AF4 | 919 | MS | 71 | 124 | Female | Black | N/A | N/A | 44 |
| 5781 | NDAR_INVDB589AU8 | 1342 | ALS | 61 | 201 | Female | White | N/A | N/A | 3 |
| 5828 | NDAR_INVUL149RBY | 1350 | Normal | 66 | 237 | Female | White | N/A | N/A | 25 |
| 5846 | NDAR_INVNR516VGD | 765 | MS | 63 | 167 | Female | White | N/A | N/A | 6 |
| 5898 | NDAR_INVAA510YXW | 1136 | ALS | 68 | 282 | Female | Asian | Negative | Negative | 17 |
| 6065 | NDAR_INVBK320LJG | 1320 | Normal | 55 | 181 | Male | White | Negative | Negative | 22 |
| 6195 | NDAR_INVEBAKXX7M | 1302 | ALS | 55 | 10 | Male | Unknown | N/A | N/A | 24 |

Note:

SID: Subject Identification number; GUID: Globally Unique Identifier; BW: Brain Weight(g); HIV: Human immunodeficiency virus; HBsAG: Hepatitis B surface antigen; N/A: not available; MS: Multiple Sclerosis; ALS: Amyotrophic Lateral Sclerosis; Normall: Unaffected Control.
